# Mechano-biological and bio-mechanical pathways in cutaneous wound healing

**DOI:** 10.1101/2022.07.28.501924

**Authors:** Marco Pensalfini, Adrián Buganza-Tepole

## Abstract

Skin injuries heal through coordinated action of fibroblast-mediated extracellular matrix (ECM) deposition, ECM remodeling, and wound contraction. Defects involving the dermis result in fibrotic scars featuring increased stiffness and altered collagen content and organization. Although computational models are crucial to unravel the underlying biochemical and biophysical mechanisms, simulations of the evolving wound biomechanics are seldom benchmarked against measurements. Here, we leverage recent quantifications of local tissue stiffness in murine wounds to refine a previously-proposed systems bio-chemo-mechanobiological finite-element model. Fibroblasts are considered as the main cell type involved in ECM remodeling and wound contraction. Tissue rebuilding is coordinated by the release and diffusion of a cytokine wave, *e*.*g*. TGF-β, itself developed in response to an earlier inflammatory signal triggered by platelet aggregation. We calibrate a model of the evolving wound biomechanics through a custom-developed hierarchical Bayesian inverse analysis. Further calibration is based on published biochemical and morphological murine wound healing data over a 21-day healing period. The calibrated model recapitulates the temporal evolution of: inflammatory signal, fibroblast infiltration, collagen buildup, and wound contraction. Moreover, it enables *in silico* hypothesis testing, which we explore by: (i) quantifying the alteration of wound contraction profiles corresponding to the measured variability in local wound stiffness; (ii) proposing alternative constitutive links connecting the dynamics of the biochemical fields to the evolving mechanical properties; (iii) discussing the plausibility of a stretch- *vs*. stiffness-mediated mechanobiological coupling. Ultimately, our model challenges the current understanding of wound biomechanics and mechanobiology, beside offering a versatile tool to explore and eventually control scar fibrosis after injury.

**Author summary:** Wounds constitute a major healthcare burden, often yielding overly stiff scars that feature altered collagen content and organization. Accurate computational models have the potential to impact the understanding, treatment, and ultimately the outcome of wound healing progression by highlighting key mechanisms of new tissue formation and providing a versatile platform for hypothesis testing. However, the description of wound biomechanics has so far been based on measurements of uninjured tissue behavior, limiting our understanding of the links between wound stiffness and healing outcome. Here, we leverage recent experimental data of the local stiffness changes during murine wound healing to inform a computational model. The calibrated model also recapitulates previously-measured biochemical and morphological aspects of wound healing. We further demonstrate the relevance of the model towards understanding scar formation by evaluating the link between local changes in tissue stiffness and overall wound contraction, as well as testing hypotheses on: (i) how local tissue stiffness is linked to composition; (ii) how a fibrotic response depends on mechanobiological cues.

## Introduction

Caused by a variety of possible conditions, including surgeries, traumas, and pathologies, wounding of the skin triggers a well-coordinated repair program that aims to rebuild the damaged tissue and recover its function via biological, chemical, and physical events [1]. Classical descriptions of healing progression consider three overlapping but distinct stages [2, 3]: inflammation, proliferation, and remodeling. *Inflammation* begins immediately after injury, when a coagulation cascade attracts platelets to the injury site [1, 3]. Their rapid aggregation in a crosslinked fibrin mesh results in a blood clot, a provisional scaffold for inflammatory cell migration [1, 3, 4]. Platelet aggregation and degranulation triggers the release of various chemokines, including platelet-derived growth factor (PDGF), vascular endothelial growth factor (VEGF), transforming growth factor beta (TGF-β), and tumor necrosis factor alpha (TNF-α) [1, 2, 4, 5]. These play a key role in recruiting neutrophils, which are the first inflammatory cells to infiltrate the wound and contribute to fight pathogens and avoid infection [2, 4]. A second wave of inflammatory cells involves monocyte migration, driven by chemoattractants such as monocyte chemotactic protein 1 (MCP-1) [2] and TGF-α [5]. Monocytes differentiate primarily into macrophages [2, 3, 5], which amplify earlier wound signals by releasing growth factors such as PDGF, VEFG, TGF-β, and fibroblast growth factors (*e*.*g*. FGF-2) [1, 2, 6]. The growth factor profiles established by macrophages coordinate tissue rebuilding during the *proliferation* phase, which occurs through the activity of various cell types [3, 7]. Keratinocytes are the first to intervene, crawling over the injured tissue in the process of epithelialization to restore the skin barrier function [2, 3]. Within angiogenesis, endothelial cells contribute to form new blood vessels [2, 5]. Fibroblasts have a key role in rebuilding the dermis — the collagen-rich layer mainly responsible for the skin structural function [8, 9] — by producing and organizing the extracellular matrix (ECM) that ultimately forms the bulk of the mature scar [3, 10], in a process stimulated by TGF-β, PDGF, and FGF-2 [5] and facilitated by cell-mediated secretion of proteolytic enzymes termed matrix metalloproteinases (MMPs) [2]. PDGF and TGF-β are also key mediators of fibroblast differentiation into myofibroblasts [5], a contractile cell phenotype that tends to approximate wound edges. Both cell types exert active stresses on the surrounding ECM and regulate collagen remodeling [2, 3], which contribute to determine the geometry and mechanical properties of the scar together with externally-applied tissue deformations [7, 11, 12]. Lastly, *remodeling* is a long-term process characterized by downregulation of overall cellular activity, cell population density, and collagen remodeling [3, 5].

Defects involving the dermis result in scars that lack the organization and full functionality of unwounded skin, exhibiting excessive stiffness, reduced strength, and permanent contracture that can persist for months or even years [10, 13–15]. This represents a significant healthcare burden, with an estimated cost per wound requiring treatment of about $4^′^000 in the US [16] and an annual management cost over £5 billion in the UK [17]. Several exogenous agents, such as medications [18], environmental conditions [19], and mechanical loading [7], may negatively affect wound healing. Fibroblast differentiation, migration, and gene expression pathways is widely acknowledged to depend on biomechanical cues [20–22]. Indeed, clinical evidence demonstrates reduction of hypertrophic scar formation via disruption of relevant mechanobiological pathways [23,24] or by modulating local tissue tension [25, 26]. Moreover, negative-pressure therapy can accelerate the healing of chronic wounds through a process involving macroscopic deformations of the wound bed [27, 28], shock waves can enhance tissue vascularization, collagen synthesis, and cell proliferation [28], and ultrasounds can stimulate granulation via tissue cavitation [28]. While these mechanotherapies can influence the course and outcome of healing, their working principles remain elusive due to insufficient understanding of the corresponding biophysical phenomena [22].

Owing to the complexity of tissue repair processes, computational models have become key tools to study the interplay of biological, chemical, and physical events, as well as to formulate and test hypotheses by providing access to quantities that are otherwise hard to determine [29]. The first computational models date back to the 1990s and mainly focused on the dynamics of cell populations, described with either ordinary [30] or partial [31] differential equations (ODEs or PDEs, respectively) or with agent based models [32]. Further developments incorporated wound contraction by imposing conservation of collagen density and linear momentum for the ECM [33], including myofibroblast contributions [34]. However, these models often featured a simplistic description of mechanics, leading to a superficial treatment of the pathways linking cell behavior to mechanical cues. We and others have been interested in incorporating detailed representations of tissue mechanics into wound healing models. Bowden *et al*. [35] proposed a purely mechanical model including tissue growth, while our most recent approaches [36–38] couple basic biochemical fields with tissue nonlinear mechanics, including permanent changes in shape and stiffness that result from growth and remodeling. Importantly, all these models adopt mechanical constitutive parameters that are representative of uninjured skin, strongly limiting their relevance towards investigating the link between ECM biomechanics and the outcome of healing.

Here, we set out to overcome this limitation by leveraging one of the very few available experimental datasets on the time-course evolution of wound mechanics at physiological deformation levels, which have been recently measured on murine tissue by Pensalfini *et al*. [39]. Through a custom-developed hierarchical Bayesian calibration procedure, we establish the change in mechanical behavior during healing, and use the calibrated constitutive model to refine our systems bio-chemo-mechanobiological finite-element (FE) model of wound healing. We further leverage the versatility of our model to test hypotheses regarding the link between tissue composition and evolving tissue stiffness, as well as the role of mechanobiological coupling to trigger fibrosis.

## Materials and methods

### Systems-mechanobiological model of wound healing

The Lagrangian FE model that we use is publicly available [40] and follows closely our original formulation [37]. Here, we briefly state the main equations and modeling assumptions, both for completeness and to reflect changes from our previous work.

### Kinematics and modeled fields

Following standard continuum mechanics notation, the current tissue geometry is described by the coordinates ***x***. The reference configuration, ***X***, coincides with the initial tissue geometry in its *ex vivo*, unloaded state. Wound healing is simulated starting from an intermediate state, ***x***^*i*.*v*.^, accounting for *in vivo* skin pre-tension. Local deformation is captured by the deformation gradient tensor, ***F*** = *∂****x****/∂****X***.

Motivated by the overview provided in the Introduction, we model the biochemical fields by grouping the release of pro-inflammatory cytokines into two waves. The first signal, *α*, is triggered upon platelet aggregation and helps direct inflammatory cells such as neutrophils and macrophages towards the wound bed. The second wave, *c*, represents the growth factors and cytokines that coordinate and regulate tissue rebuilding and remodeling via fibroblasts and myofibroblasts. Being mainly interested in new tissue formation and remodeling, we limit our description of inflammation to the fields *α* and *c*, avoiding explicit modeling of the corresponding cell species. Accordingly, the cell population density, *ρ*, coincides with the amount of fibroblasts/myofibroblasts in the tissue, owing to their widely-recognized role towards determining ECM deposition and organization in wounds and scars. Lastly, the tissue composition is mainly described by its collagen content, *ϕ*_*c*_, and by the plastic deformation, ***F***^*p*^, which reflects on the permanent stretch ratios, 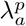 and 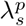, measured along the in-plane eigenvectors of ***F***^*p*^, ***a***_0_ and ***s***_0_. Note that we take the values of *c, ρ, ϕ*_*c*_, 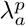, and 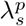 in an unwounded tissue subjected to a physiological deformation level to be 1, while *α* = 0 in such conditions.

### Balance laws for mass and linear momentum

The balance of linear momentum follows the standard relation ∇ · ***σ*** = **0**, where ***σ*** denotes the Cauchy stress tensor and is determined by two contributions:

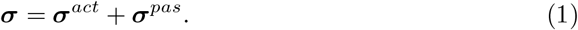

In Eq. (1), ***σ***^*act*^ is the active stress exerted by the cell population, *ρ*, on the collagenous ECM, *ϕ*_*c*_, and our model assumes that it can be influenced by two factors, *cf*. Eqs. (10,11) and Fig. 1: (i) the mechanical state of the wound, affected by the ECM deformation and its material properties; (ii) the cytokines *c*, affecting cell-mediated ECM contraction. The passive stress, ***σ***^*pas*^, only depends on the mechanical state of the tissue via Eq. (2), *cf*. Fig. 1.

**Figure 1.**
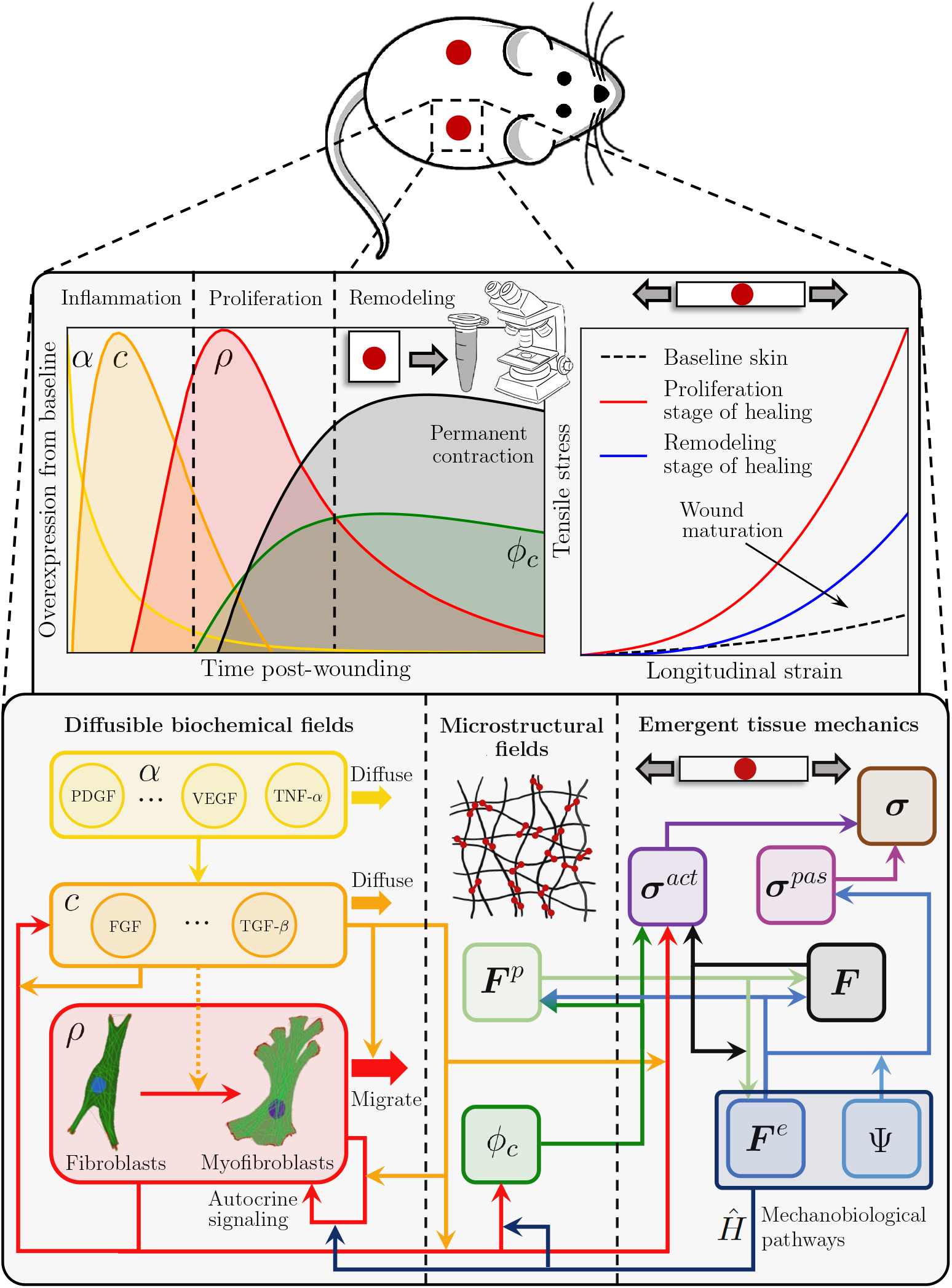
Schematic summarizing the developed systems-mechanobiological model of wound healing, which aims to capture the temporal evolution of key biochemical, microstructural, and macroscopic mechanical and geometrical variables by representing the cell and tissue regulatory pathways and their interaction across structural scales.

The balance of mass for the fields *α, c*, and *ρ* also follows standard equations. For instance, the second inflammatory signal must satisfy the relation *ċ* + ∇ · ***q***_*c*_ = *s*_*c*_, where *s*_*c*_ and ***q***_*c*_ are the source and flux terms in the current configuration, respectively. In the Lagrangian setting, the flux of *c* is expressed by ***Q***_*c*_ = *J****F***^−1^***q***_***c***_. Conversely, the tissue microstructural fields, 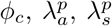, are updated locally and without accounting for any diffusion; their rate of change is defined later on, along with the corresponding constitutive equations. Owing to lack of diffusion in the microstructural fields, we refer to *α, c*, and *ρ* as *diffusible* biochemical fields (Fig. 1).

### Constitutive equations for the tissue mechanical behavior

The passive stress in the tissue derives purely from the elastic part of the deformation gradient tensor ***F***^*e*^ = ***FF***^*p*−1^ and is assumed to follow from a hyperelastic potential similar to the one used in the Gasser-Ogden-Holzapfel (GOH) model [41]:

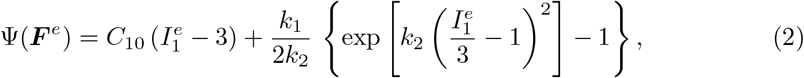

where 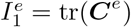 is the first invariant of the elastic right Cauchy-Green deformation tensor, ***C***^*e*^ = ***F***^*e*⊤^***F***^*e*^, and *C*_10_, *k*_1_, *k*_2_ are material parameters. This constitutive model describes the characteristic J-shaped stress-strain response observed in many soft biological tissues [41–46] by assuming a neo-Hookean ground substance with shear modulus *μ*_0_ = 2*C*_10_ and a collagen-based network whose emergent stiffening upon stretching is controlled by the phenomenological parameters *k*_1_ and *k*_2_. Note that we consider a simplified model with respect to the original formulation [41], by assuming that skin and wound/scar tissues are isotropic materials subjected to plane stress conditions, since the modeled skin region has thickness much smaller than its in-plane dimensions. The constitutive parameters are first determined from experimental data [39] in order to perform wide-range model calibration, and later derived from constitutive equations linking them to the time-course evolution of the microstructural fields.

### Constitutive equations for the diffusible biochemical fields

The fluxes of *α, c*, and *ρ* in the reference configuration are expressed by:

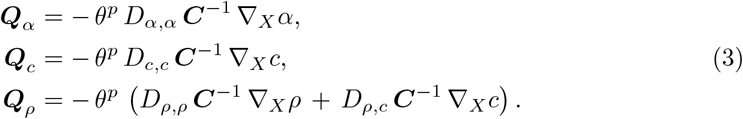

Since inflammatory cell recruitment is mostly completed within the first few days after injury [5], we assume an exponentially-decaying source term for *α*, with rate *d*_*α*_:

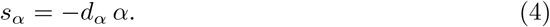

The source term for the second inflammatory wave, *c*, accounts for its dependence on the first inflammatory wave, *α*, and on the fibroblast/myofibroblast population density, *ρ*:

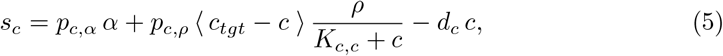

where *c*_*tgt*_ is an attractor for *c* and is selected to ensure homeostasis of this biochemical field for an unwounded tissue subjected to physiological deformation, while ⟨ ⟩ denote the Macauley brackets that prevent the second term on the right-hand side of Eq. (5) from becoming negative when *c* > *c*_*tgt*_. *K*_*c,c*_ determines the saturation of *c* in response to itself, *p*_*c,α*_ and *p*_*c,ρ*_ are coefficients capturing the effects of *α* and *ρ* on the release of cytokines, *c*, (Fig. 1), and *d*_*c*_ is the decay rate when all production terms are zero.

The source term for *ρ* is an extension of logistic models with cytokine feedback and mechanobiological coupling, and is purely a reformulation of our previous works [37, 38]:

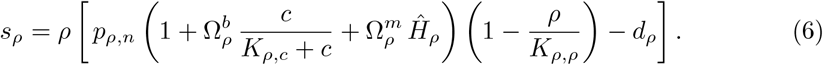

The function *Ĥ*_*ρ*_ encodes the dependence of fibroblast/myofibroblast proliferation on the mechanical state of the tissue (Fig. 1), as discussed in more detail below. *p*_*ρ,n*_ defines the natural mitotic rate of the cells in the absence of cytokines and mechanical effects (*c* = 0, *Ĥ*_*ρ*_ = 0), while 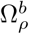 and 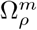 are coefficients capturing the enhanced cell proliferation in response to *c* and *Ĥ*_*ρ*_ (Fig. 1). *K*_*ρ,c*_ and *K*_*ρ,ρ*_ determine the saturation of *ρ* in response to *c* and to itself, and *d*_*ρ*_ is again the decay rate when all production terms are set to zero. We select *d*_*ρ*_ to ensure that *ρ* respects homeostasis for an unwounded tissue subjected to physiological deformation.

### Constitutive equations for the tissue microstructural fields

The tissue microstructure changes in two ways during wound healing: one of them is the change in composition, *e*.*g*. the collagen mass fraction *ϕ*_*c*_, and the other is the change in permanent deformation, ***F***^*p*^, which reflects on the permanent stretch ratios, 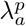 and 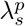.

Similar to our previous works [37, 38], we assume that *ϕ*_*c*_ depends linearly on the fibroblast/myofibroblast population density, *ρ*, in a way that is mediated by *c* (Fig. 1):

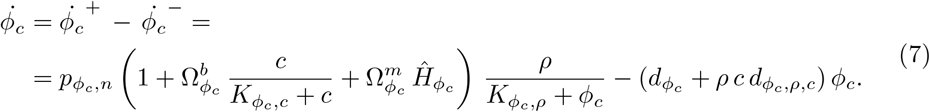

The function 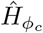 encodes the dependence of cell-mediated collagen deposition on the mechanical state of the tissue (Fig. 1). For simplicity, we set 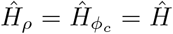, *i*.*e*. we assume that mechanical cues impact cell proliferation and collagen deposition in the same fashion up to a scaling factor, 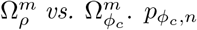 defines the natural rate of collagen deposition in the absence of cytokines and mechanical effects (*c* = 0, 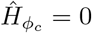), while 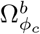 and 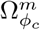 are coefficients capturing the enhanced collagen deposition in response to *ρ* and 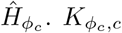 and 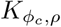 determine the saturation of *ϕ*_*c*_ in response to *c* and to *ρ*. Note that, beside a spontaneous decay mediated by the coefficient 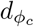, selected to ensure homeostasis of *ϕ*_*c*_ in unwounded physiological conditions, the collagen degradation rate 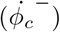 also depends on *ρ* and *c* via 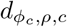, capturing the role of cells within MMP production [2] and the corresponding modulation by cytokines [47].

Lastly, the remodeling law operates independently along the principal directions ***a***_0_ and ***s***_0_ according to the equation (Fig. 1):

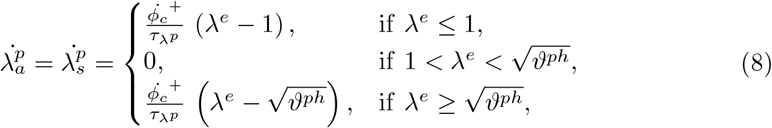

where *λ*^*e*^ is the current elastic stretch of the tissue along the direction of interest, 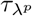 is the time constant for tissue growth in either direction, and *ϑ*^*ph*^ captures the elastic areal deformation of the tissue in its physiological *in vivo* state. Different from our previous approaches [36–38], Eq. (8) implies that the tissue grows when stretched past its physiological state, shrinks when subjected to prolonged compression, but accumulates no permanent deformation when stretched to sub-physiological levels.

### Mechano-biological and bio-mechanical pathways

The modeled biochemical fields can determine (bio→mechanics) and be determined (mechano→biology) by the mechanical state of the tissue ECM in multiple ways (Fig. 1). A first relevant bio-mechanical pathway has been discussed in the previous section: plastic deformation is influenced by collagen turnover, itself a function of cell activity. Plastic deformation also dissipates elastic energy, affecting tissue deformation, ***F***^*e*^. This has a direct mechano-biological effect on the cell population and collagen deposition, as mediated by the function *Ĥ, cf*. Eqs. (6, 7). A leading hypothesis in the field, which we have also used previously [37, 38], postulates a dependence of *Ĥ* on strain:

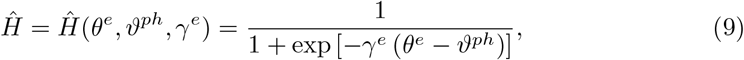

where *θ*^*e*^ = ||cof***F***^*e*^ · **N**|| captures the current in-plane elastic tissue deformation, while *γ*^*e*^ controls the slope of *Ĥ* around its midpoint, *ϑ*^*ph*^. While the majority of this manuscript adopts Eq. (9), we will also explore an alternative coupling as part of our hypothesis testing efforts, *cf*. Results.

A further bio-mechanical pathway of interest is represented by cell-induced contraction, yielding an active tension that depends on *ρ* and *c* (Fig. 1):

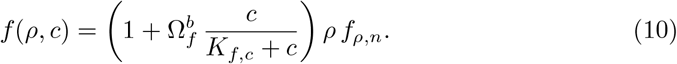

In the above expression, which is purely a reformulation of what we have previously used [37, 38], *f*_*ρ,n*_ is the baseline tension exerted by a physiological population of fibroblasts/myofibroblasts on the surrounding ECM in the absence of any cytokine (*ρ* = 1, *c* = 0), 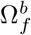 captures the tension increase in response to *c, e*.*g*. due to enhanced fibroblast-myofibroblast transition, and *K*_*f,c*_ determines the saturation of *f* in response to *c*. The active stress resulting from the tension introduced in Eq. (10) reads:

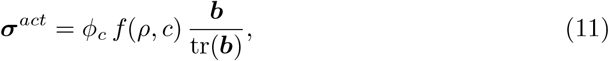

where ***b*** = ***FF***^⊤^. Note that, owing to the assumption of tissue isotropy, the current expression for active stress is simplified with respect to our previous approach [37], which adopted a deformed structural tensor based on a direction of anisotropy.

### Hierarchical Bayesian calibration of mechanical parameters

To determine the constitutive parameters of unwounded and wounded tissues at various healing time points, we leverage the experimental measurements previously presented in Pensalfini *et al*. [39], where several specimens including a 7- or 14-day-old wound were subjected to *ex vivo* uniaxial tensile tests and compared to the mechanical response of unwounded skin. Contrary to most traditional material parameter-fitting approaches, which either focus on the average measured mechanical response for a set of homogeneous specimens [45, 48, 49], or treat each tested specimen completely independently [43, 45, 50, 51], we adopt a hierarchical Bayesian calibration procedure.

Consider a generic mechanical constitutive model that can be specified by prescribing an *m*-tuple of parameters, *μ* = (*μ*_1_, …, *μ*_*m*_), providing a deterministic relation between applied deformation, *e*.*g*. the stretch ratio *λ*, and stress: *P* = *P* (*μ, λ*). We wish to determine values of *μ* yielding the experimentally-measured stresses for each of the *N*_*s*_ tested tissue specimens at each healing time point 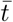. We denote each such parameter set as 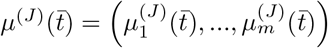, with *J* = 1, …, *N*. Since wound infliction and healing progression are presumably the major factors determining the mechanical differences measured in Ref. [39], we reason that all the *N*_*s*_ parameters 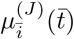, where 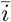 denotes a fixed value of *i*, must share some underlying similarity. Indeed, all 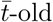 tested wounds were obtained from mice of comparable genotype, age, sex, *etc*… and subjected to similar handling until the moment of tissue excision and testing, which were also performed using the same devices and data analysis procedures. This suggests that all the 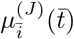 can be regarded as originating from a common probability distribution; similar considerations apply to the unwounded skin specimens. To restrict the mechanical constitutive parameters to be non-negative, we assume that 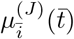 originate from a log-normal distribution,

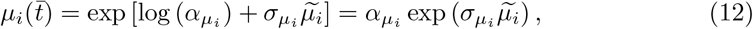

where 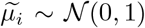 is a normally-distributed variable (𝒩) with zero mean and unit variance, while 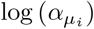 and 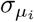 are the expected value and standard deviation of 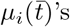 natural logarithm. Importantly, 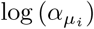 and 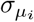 are not fixed values, but they also originate from distributions (Fig. 2), highlighting the *hierarchical* (or nested, or multilevel) structure of the posed statistical model, where the moments of the mechanical parameter distributions are themselves obtained from distributions. To stress the generating role of 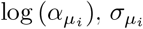, and 𝒩(0, 1) with respect to 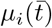, we refer to the former as *hyper* distributions. In the absence of more detailed information, we will assume uniform hyperdistributions (𝒰) with lower and upper bounds depending on the subscript 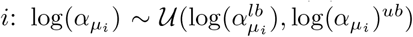 and 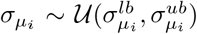. Thus, each independent sampling of the hyperdistributions 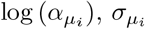, and 𝒩(0, 1) generates, via Eq. (12), one value for the *i*-th entry of the *m*-tuple of mechanical parameters, 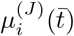. In turn, the mechanical constitutive model provides a deterministic relation between stretch and stress, 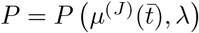 (Fig. 2), and the experimental me asurem ent process introduces further uncertainty, which we model as Gaussian noise 𝒩 (*P*, Σ^2^).

**Figure 2.**
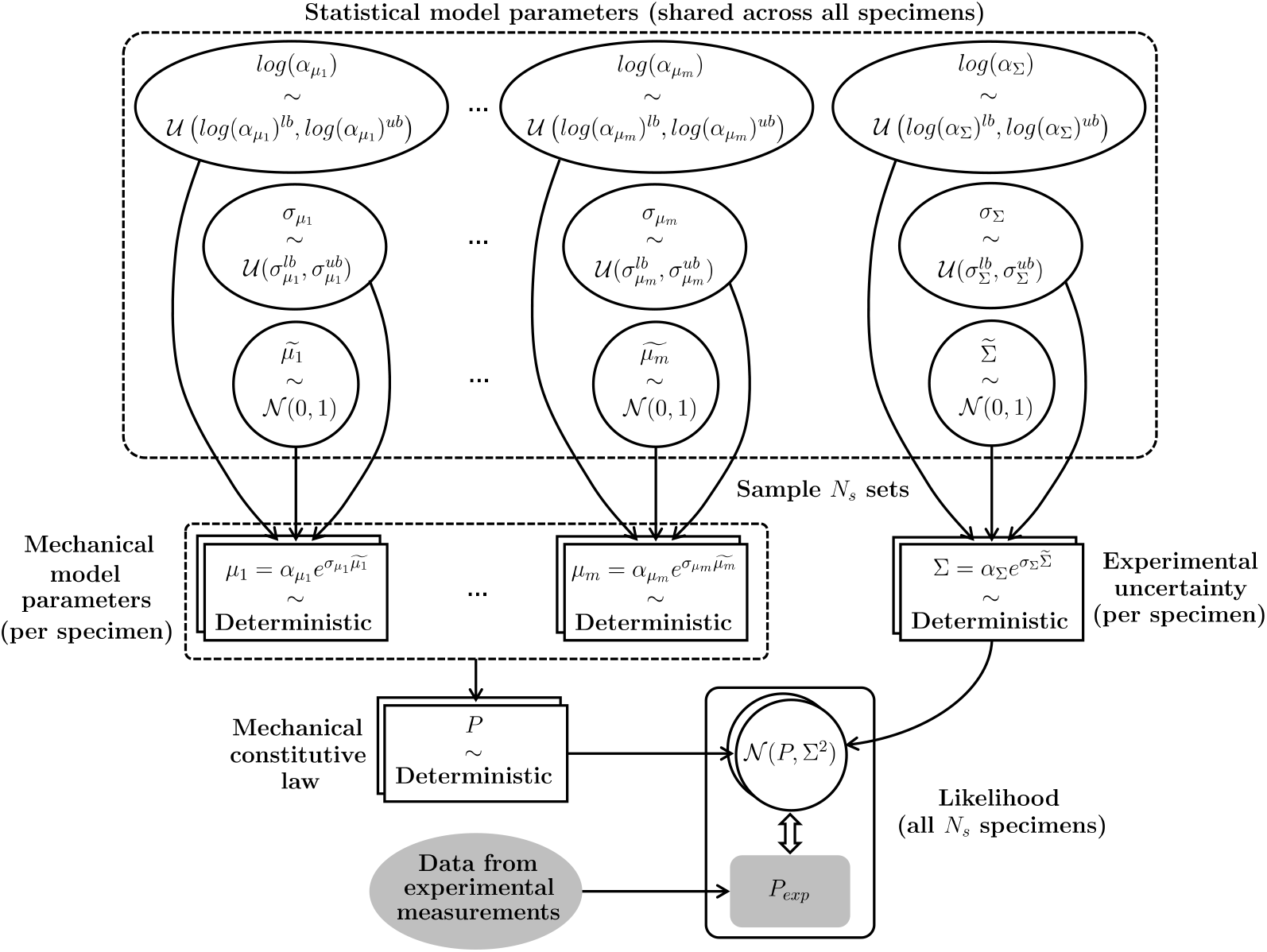
Schematic of the hierarchical Bayesian model posed to capture the experimentally-measured mechanical response of a set of *N*_*s*_ inter-related tissue specimens. A set of 3 (*m* + 1) hyperdistributions, common across all specimens, generates the *mN*_*s*_ mechanical model parameters corresponding to each tested specimen, and the *N*_*s*_ parameters representing experimental uncertainty. These parameters yield deterministic predictions for the mechanical behavior of each specimen, which are to be compared to the corresponding experimental evidence in order to establish a best-fitting set of hyperparameters.

While the above specifications define the *direct* link between a set of statistical model parameters, 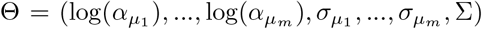, and the measured tensile response predicted by the model, 𝒩(*P*, Σ^2^), we are interested in the corresponding *inverse* relation, from the experimental measurements to the parameters, Θ. In Bayesian terms, the hierarchical model prescribes the *likelihood* p (𝒩(*P*, Σ^2^) | Θ), *i*.*e*. the probability distribution of the observed data given a set of parameters, and we wish to determine the *posterior* p (Θ | *P*_*exp*_), *i*.*e*. the probability distribution of the parameters given the data, *P*_*exp*_. According to Bayes’ theorem,

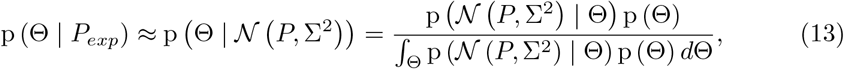

where p (Θ) is the *prior, i*.*e*. the probability distribution of the parameters Θ before any observation has been made, and ∫_Θ_ p (𝒩 (*P*, Σ^2^) | Θ) p (Θ) *d*Θ is the *evidence*, a normalization factor coinciding with the probability distribution of the observed data independently from any parameter set. To determine the posterior without computing the evidence, whose integral can easily become intractable, it is possible to resort to numerical methods such as Markov Chain Monte Carlo (MCMC) or Variational Inference (VI) [52]. A key difference between the two approaches is that MCMC assumes no model for the posterior, while VI casts inference as an optimization problem and seeks the best approximant of the posterior within a parameterized family of distributions according to a suitable cost function, *e*.*g*. the Kullback-Leibler divergence, or a likelihood function, *e*.*g*. the Evidence Lower Bound (ELBO) [52]. This allows reducing the variance of the method at the cost of introducing some bias, such that VI approaches are generally less accurate than MCMC but tend to be faster and scale better to large datasets [52]. Given the complexity of our hierarchical model and the fairly large datasets that we aim to fit, featuring thousands of experimental data points, we adopt a VI approach and specify the model using the Python-based probabilistic programming framework PyMC3 [53]. Specifically, we adopt a gradient-based approach known as Automated Differential Variational Inference (ADVI) [54] and assume that the posterior follows a spherical Gaussian distribution without correlation of parameters, which we estimate by maximizing the ELBO [53]. The corresponding code is publicly available [40].

## Results

### Evolution of wound mechanical behavior throughout healing

To determine GOH constitutive parameters describing the tensile experiments performed in Ref. [39], we focus on each of the three available datasets and conduct separate parameter optimizations for the unwounded skin specimens (*N*_*s*_ = 8), the 7-day-old wounds (*N*_*s*_ = 8), and the 14-day-old wounds (*N*_*s*_ = 8). For simplicity of analysis, we focus on the wound core regions identified in Ref. [39] and assume that they were subjected to uniaxial tensile loading, neglecting any possible influence of the surrounding tissue on the measured response. We also assume a tissue thickness of 1.7 mm, in line with [10].

For each of the three separate parameter optimizations, we train the hierarchical model with 200^′^000 samples by prescribing fairly broad search ranges, *cf*. S1 Table, and ensuring convergence of the ELBO, *cf*. S1 Fig. We then use the calibrated statistical models to generate 10^′^000 independent samples, each of them yielding *N*_*s*_ = 8 realizations of the mechanical parameters *C*_10_, *k*_1_, and *k*_2_, one for each specimen in the considered dataset. This allows visualizing the specimen-specific mechanical parameter posteriors, which are reported in S2 Fig, S3 Fig, and S4 Fig, along with the priors, traces, and posteriors of the corresponding statistical model parameters. For simplicity, Fig. 3a-c only shows the median values of the specimen-specific mechanical parameter posteriors (dots), along with boxplots indicating the corresponding median (orange line), interquartile range (box), and the 95% confidence interval (CI, indicated by the extension of the whiskers) for each of the three analyzed datasets. Remarkably, the posterior of the constitutive parameter *k*_2_, which mainly controls tissue stiffening at large stretches, can only be inferred for the unwounded specimens (Fig. 3c and S2 Fig), which are indeed those consistently reaching the largest deformations (Fig. 3d *vs*. Fig. 3e,f). Instead, for the wounded specimens, we fix *k*_2_ = 0.88 according to the median of the specimen-specific posteriors obtained for unwounded skin (Fig. 3c) and restrict our analysis to the parameters *C*_10_ and *k*_1_.

**Figure 3.**
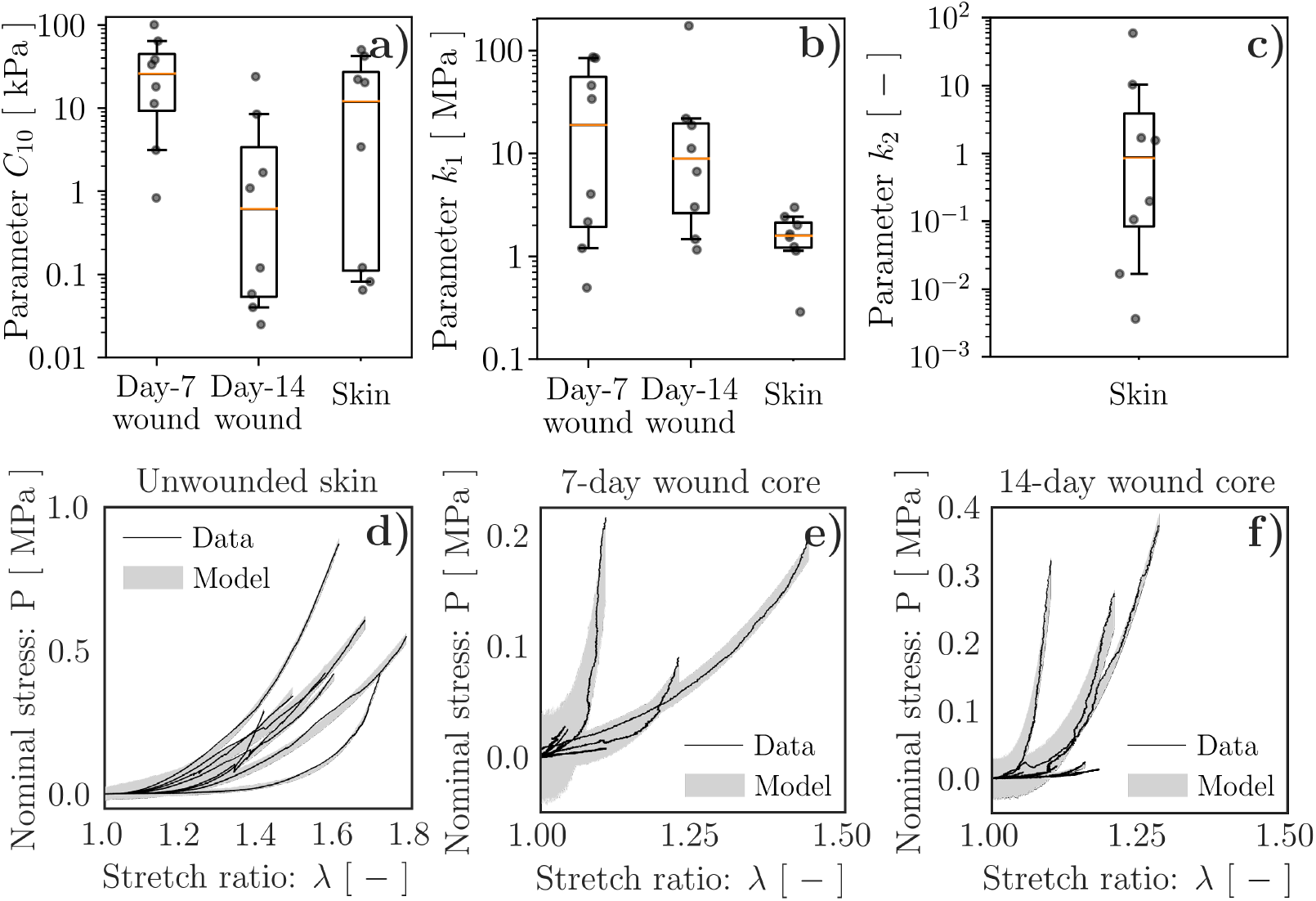
Hierarchical Bayesian calibration of tissue mechanical parameters. (**a**) *C*_10_, corresponding to the behavior of the non-collagenous ground substance, exhibits marked decrease between 7- and 14-days post-wounding, while mostly remaining within the broad range of values characterizing unwounded skin. (**b**) *k*_1_, corresponding to the behavior of the tissue collagenous matrix, tends to decrease between 7- and 14-days post-wounding, but is typically larger in the wounds than in unwounded skin. (**c**) *k*_2_, relating to the large deformation behavior of the tissue collagenous matrix, can only be inferred for unwounded skin due to the limited deformability of wounded tissues prior to failure. (**d**–**f**) Model-based predictions of the specimen tensile behavior accounting for the experimental uncertainty match the experimental data reasonably well. The dots in (**a**–**c**) indicate median values of the specimen-specific mechanical parameter posteriors, cf. S2 Fig, S3 Fig, and S4 Fig. The boxplots in (**a**–**c**) are constructed based the values indicated by the dots, with orange lines denoting the median and the extension of the whiskers denoting the 95% CI. The shadings (**d**–**f**) indicate the 95% CI obtained from 1^′^000 random tensile curves generated using the calibrated Bayesian model and accounting for experimental uncertainty.

Despite the large variability in the inferred mechanical parameters, which is certainly not unexpected when quantifying the properties of biological materials, the adopted hierarchical model provides information on each tested specimen, allowing us to discuss the evolution of *C*_10_ and *k*_1_ throughout healing. On the one hand, *C*_10_, whose median value across 7-day-old specimens is about 2.2× the one of unwounded skin, reaches about 1*/*20 of the unwounded value at day 14, showing a 42.5× reduction. Instead, *k*_1_ is consistently larger in the wounds (11.9× at day 7, 5.6× at day 14) than in the unwounded skin specimens, despite a 2.1× reduction between days 7 and 14. Since the GOH model uses *k*_1_ to capture the stiffening of the collagenous ECM, the evolution of this parameter can be interpreted as indicative of pronounced and sustained tissue fibrosis in the wound/scar with respect to the unwounded tissue. Concomitantly, the GOH model parameter C_10_, capturing the mechanical contribution of the non-collagenous ground substance, also appears significantly affected by wound healing progression, suggesting marked softening of this tissue component.

Lastly, we access the calibrated statistical model traces to visualize the tensile curve posteriors. For each specimen, we randomly generate 1^′^000 curves that account for experimental uncertainty, as captured by the modeling parameter Σ, and confirm that the 95% CI of the model-based predictions match the experimental measurements (Fig. 3d–f). Thus, the determined constitutive parameters capture the tensile response of the tissue specimens, suggesting that they can be used to infer the time-evolving mechanical behavior of wounded skin throughout healing.

### Influence of wound deformability on the healing outcome

Having established plausible ranges and trends for the mechanical constitutive parameters of wounded and unwounded tissues, we aim to investigate their influence on the evolution and outcome of healing. To this end, we first perform wide-range calibration of our custom FE model using the median values of the constitutive parameters at each time point, and then vary them according to the determined 95% CI.

To simulate *in vivo* wound healing, we start from a square skin patch with side length 50 mm in its reference state (Fig. 4a and S1 Video), and set its mechanical constitutive parameters according to the median values obtained from Bayesian calibration: 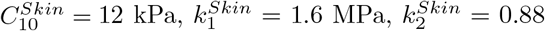. To limit the computational cost, we restrict our model to 1*/*4 of the considered patch and impose symmetric boundary conditions on *x* = 0 and *y* = 0 (*cf*. Fig. 4 and S1 Video). We approximate *in vivo* pre-tension by subjecting the unwounded tissue to an equibiaxial stretch with *λ*_*x*_ = *λ*_*y*_ = 1.15 (Fig. 4b and S1 Video), which is within the broad range of previously-reported post-excisional skin shrinkage values [51,55,56]. Following equilibration, we introduce a wound by setting the constitutive parameters *C*_10_ and *k*_1_ in a 5 mm-diameter tissue region located at the center of the computational domain to extremely small values (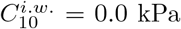 and 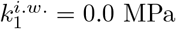, *i*.*w*.: immediately after wounding), while leaving *k*_2_ unchanged. Owing to the stark contrast between the mechanical properties of the wound and those of the surrounding skin, this causes the wound region to expand (inset in Fig. 4c and S1 Video), much like the classical problem of a membrane featuring a circular hole and subjected to tension. Shortly after infliction, at time point *d*0 (day 0 of the healing time-course, Fig. 4c), we also impose that the wound exhibits a peak in the first inflammatory wave (*α*^*d*0^ = 1), which is associated with platelet aggregation, and negligible values for the second inflammatory wave (*c*^*d*0^ = 0), cell density (*ρ*^*d*0^ = 0), and collagen content 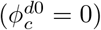. Meanwhile, all biochemical and microstructural variables in the surrounding skin have physiological values: *α*^*Skin,d*0^ = 0, *ρ*^*Skin,d*0^ = 1, *c*^*Skin,d*0^ = 1, and 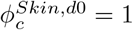. As for the mechanical parameters, we reason that no collagen deposition can occur prior to day 0. Thus, tissue integrity must be supported by the fibrin clot, leading us to assume 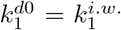 and to linearly extrapolate the value of 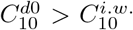 based on its values at days 7 and 14 post-wounding (Fig. 3a,b).

**Figure 4.**
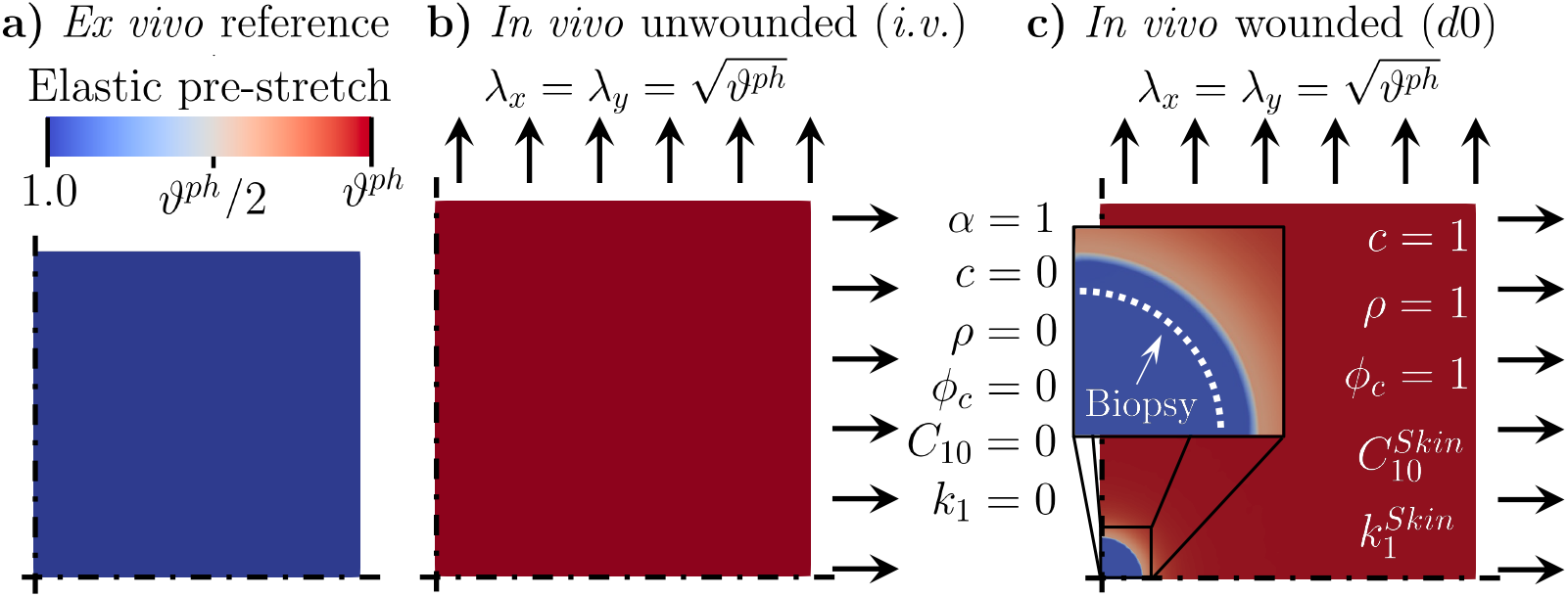
Model preparation before solving the wound healing problem. (**a**) Modeled square tissue patch, with symmetric boundary conditions along *x* = 0 and *y* = 0 and mechanical constitutive parameters corresponding to unwounded skin. (**b**) Unwounded skin patch in its *in vivo* state (*i*.*v*.), characterized by an equi-biaxial pre-stretch that can be fully released upon unloading. (**c**) Wound infliction *in vivo*, obtained by setting the mechanical parameters in a circular tissue region to extremely small values; the values of the biochemical and microstructural quantities (*α, c, ρ, ϕ*_*c*_) are also adjusted to reflect a freshly-wounded tissue. Note that, immediately after infliction, the wound enlarges due to the corresponding release of tissue pre-stretch, as shown in the inset (white dashed line *vs*. boundary of the blue region). The reached deformation is made permanent to ensure that the newly-deposited tissue has no initial stress.

We then simulate wound healing over a 21-day period by assuming that the material parameters vary linearly between their known values at days 7 and 14, and then remain constant between days 14 and 21 (Fig. 5a,b, S1 Video, and S5 Table). All other model parameters are set according to the values reported in the Supporting information (S2 Table–S5 Table) in order to match available literature data on the amount of cytokines [57–60], cells [57, 61], and collagen [62–65] in murine wounds, *cf*. Fig. 5d–f and S1 Appendix, as well as previously published data on changes in the visible wound area [10], *cf*. Fig. 5i. Lastly, the parameters of Eq. (9) are selected in order to capture previously-reported information on the stretch-dependence of human patellar tendon fibroblasts proliferation [66], *cf*. S1 Appendix.

**Figure 5.**
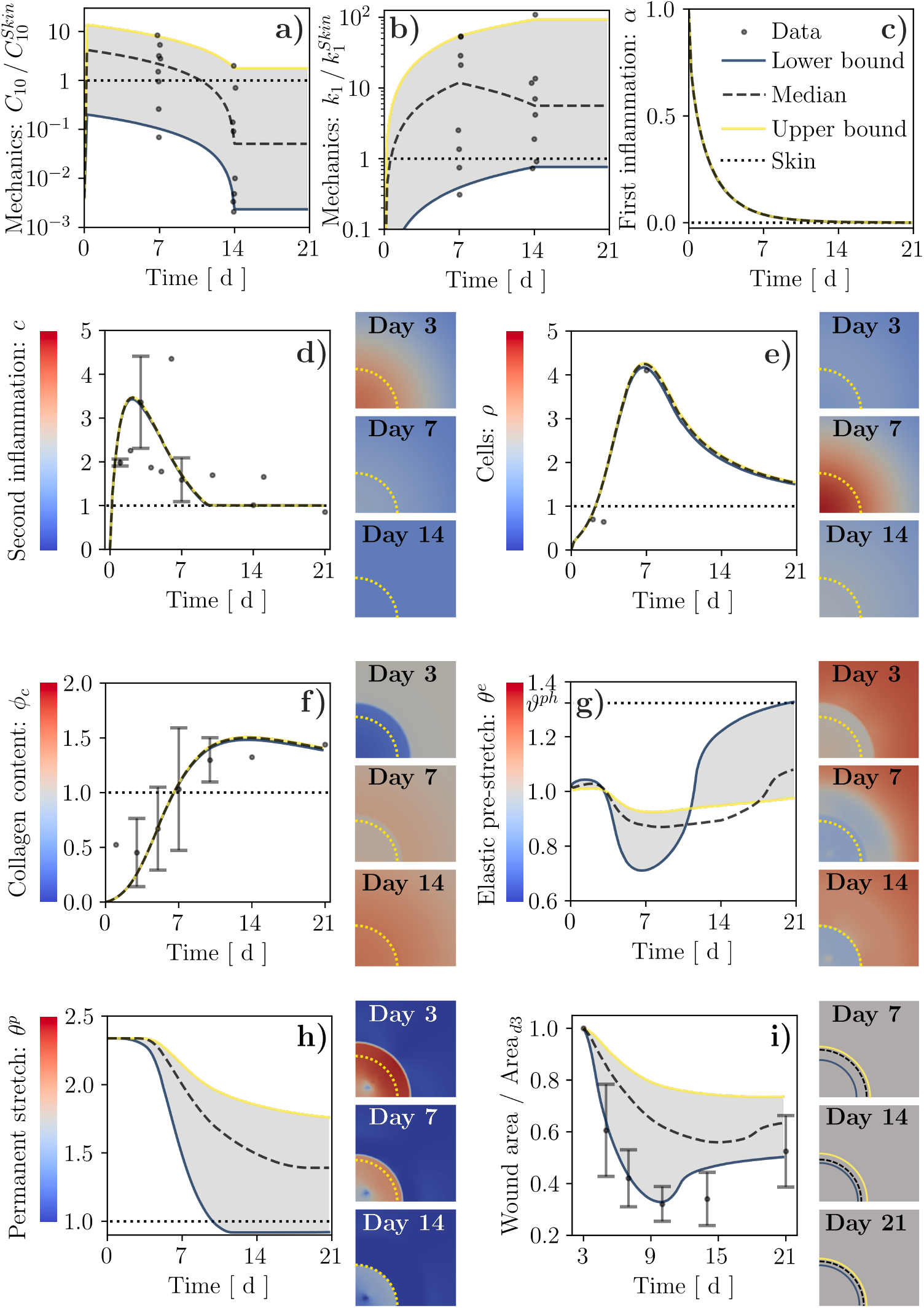
Results of wound healing simulations over a 21-day period using the wound mechanical parameters directly obtained from the Bayesian calibration procedure (median and 95% CI) and assuming linear variation between known values. (**a, b**) Hard-coded time evolution of the mechanical parameters 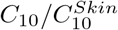 and 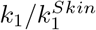, along with the corresponding values from Bayesian calibration (dots, *cf*. Fig. 3). (**c**) Decay of the first inflammatory signal, *α*, in the wound. (**d**–**h**) Time and spatial evolution of: second inflammatory signal, *c*; cell population, *ρ*; tissue collagen content, *ϕ*_*c*_; tissue elastic stretch, *θ*^*e*^; tissue plastic stretch, *θ*^*p*^. (**i**) Time evolution and illustration of wound area changes. Dots and error bars in (**d**–**i**): mean ± standard deviation of previously-published experimental data, *cf*. S1 Appendix and Ref. [10].

Irrespective of the imposed constitutive parameters, *α* simply decays exponentially to zero over 7–10 days (Fig. 5c and S1 Video) as prescribed by Eq. (4). Similarly, *c* increases from its initial value of 0 to a maximum of about 3.5 × at day 2–3 post-wounding, before returning to its physiological value of 1 in a way that is also largely independent of mechanics (Fig. 5d and S1 Video). The contours to the right of the chart in Fig. 5d show the spatial variation of *c* over time. As expected from the dependence of *c* on *α* (Eq. (5)), and based on the role of diffusion, the profiles for *c* exhibit a peak at the center of the wound in the early stages of healing (days 0–3), which diffuses smoothly into the surrounding tissue over time (days 3–7).

The cell density, *ρ*, exhibits a peak of about 4.2× around day 7 post-wounding, followed by gradual decay over time (Fig. 5e and S1 Video). The entire evolution of *ρ* lags behind that of *c*, as also visible in the corresponding contour plots. This delay originates from both fibroblast chemotaxis (Eq. (3)) and increased proliferation (Eq. (6)) in response to *c*. Fibroblasts infiltrating the wound have a key role in depositing collagen, one of the main microstructural fields in the current model, leading its content to gradually increase starting from day 7 post-wounding. Remarkably, the collagen content in the wound peaks at a value of 1.4–1.5×, which is first reached around day 10 and persists until day 21 (Fig. 5f and S1 Video). Note that, while *ρ* and *ϕ*_*c*_ explicitly depend on the mechanical deformation of the tissue (Eqs. (6,7)), varying the mechanical constitutive parameters according to the determined 95% CI has almost no influence on Fig. 5e,f. This counterintuitive finding will be discussed in a later section.

The other main microstructural field that we quantify from the FE simulations is the plastic deformation, 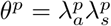 (Fig. 5h), which is governed by Eq. (8) and is intimately related to the elastic deformation, *θ*^*e*^ (Fig. 5g). Shortly after injury, the wound void — enlarged due to *in vivo* pre-tension — is filled by a fibrin clot made of a virgin material that has no initial stress, as ensured by setting its plastic deformation to an initial value *θ*^*p*^ = *θ*^*e, i*.*w*.^, where *θ*^*e, i*.*w*.^ ≈ 2.3 is the elastic deformation of the wound void after it has enlarged as a consequence of wound infliction, measured with respect to the reference configuration, ***X*** (S1 Video). Correspondingly, the elastic deformation in the clot is initially 1 and its value throughout healing is affected both by the active stress applied by the infiltrating fibroblasts and by the growth/shrinkage of the wound tissue (Eq. (8)). Notably, *θ*^*e*^ < 1 for the majority of wound healing progression, determining a progressive decrease of the plastic deformation throughout healing (Fig. 5h and S1 Video). Correspondingly, the visible wound area (Fig. 5i) decreases drastically over the first 10 days post-wounding, before stabilizing or even slightly increasing towards the end of the considered 21-day period. Note that, unlike the amount of elastic or plastic deformation in the wound, its size is typically measured in wound healing experiments, allowing our model to be compared to quantitative data such as those presented in Ref. [10].

Remarkably, the temporal evolution of the fields directly associated with deformation is strongly influenced by the constitutive behavior of the wound ECM. Both the plastic deformation and wound area are smaller/larger for softer/stiffer ECMs, while the influence of *C*_10_ and *k*_1_ on the elastic deformation is more complex and depends on the ratio between the ECM deformability in the wound and that of the surrounding skin. Indeed, when the wound ECM is softer than the surrounding skin for the majority of the healing time-course (lower bound curves in Fig. 5), the infiltrating cells can easily contract the wound, resulting in a strong initial decrease in *θ*^*e*^ — until a minimum of about 0.7 around day 7 — that yields a drastic reduction in *θ*^*p*^, as prescribed by Eq. (8). Around day 14, the plastic deformation has practically vanished in such case, so that the active stress reduction associated with the downregulation of *c* and *ρ* allows the tissue to approach its physiological elastic deformation, *ϑ*^*ph*^. Conversely, when the wound ECM is consistently stiffer than the surrounding tissue (median and upper bound curves in Fig. 4), the effect of the active stresses on the wound deformation is mitigated, resulting in a more modest reduction of *θ*^*p*^ during the early stages of healing, which never approaches the value of 1. This leads to a much more modest increase in wound deformation when the active stresses are subsequently reduced, such that *θ*^*e*^ never approaches *ϑ*^*ph*^ in this case.

The present wide-range model calibration allows recapitulating the temporal evolution of several key aspects of wound healing, such as infiltration of cytokines and fibroblasts, collagen deposition, as well as the size and deformation of a developing scar. As such, our model offers a versatile platform to address the plausibility of alternative hypotheses concerning the biomechanical and mechanobiological pathways involved in wound healing. A first example is provided by the present analysis, demonstrating that wound ECM deformability can have a major influence on the healing outcome by significantly affecting the wound contraction profiles.

### Linking the wound mechanical behavior to tissue microstructure

Having established a reliable model of wound progression, we now aim to propose plausible links between the emergent mechanical behavior of the healing tissue and the microstructural fields that, in turn, depend on the biochemical fields. Hence, we turn our attention to replacing the hard-coded evolution of wound mechanical properties by constitutive hypotheses. Collagen being among the major determinants of soft tissue biomechanics [8], a common approach in the literature [67–70] — which we have also previously followed [37, 38] — is to make *k*_1_ proportional to *ϕ*_*c*_. In addition, our material parameter calibration indicated a clear reduction of the value of *C*_10_, capturing the mechanical behavior of the tissue non-collagenous ground substance, throughout healing (Fig. 3a). Accordingly, we posit that *C*_10_ might represent the mechanical contribution of the fibrin clot that is formed at the onset of the healing response and is gradually depleted by the infiltrating cells via fibrinolytic enzymes and MMPs [5]. Thus, we introduce a microstructural field encoding the wound fibrin content, 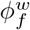, and assume a purely decaying temporal evolution mimicking that of collagen:

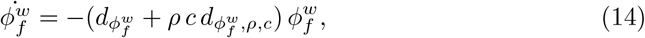

where 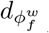 defines the extent of spontaneous fibrin decay and 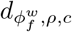 defines the magnitude of cell-mediated fibrin depletion, which we assume to be affected by *c* in line with the overall modulation of cell activity by cytokines. Note that the microstructural field 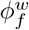 is specific to the wound, hence the superscript *w*. Here, we set 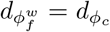 and choose 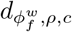 to ensure that most fibrin decays prior to day 7 (Fig. 6c). Similar to the classical link between *k*_1_ and *ϕ*_*c*_, we also assume proportionality of 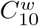 to 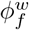, leading to the following relations between tissue mechanics and microstructure:

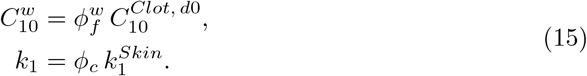

**Figure 6.**
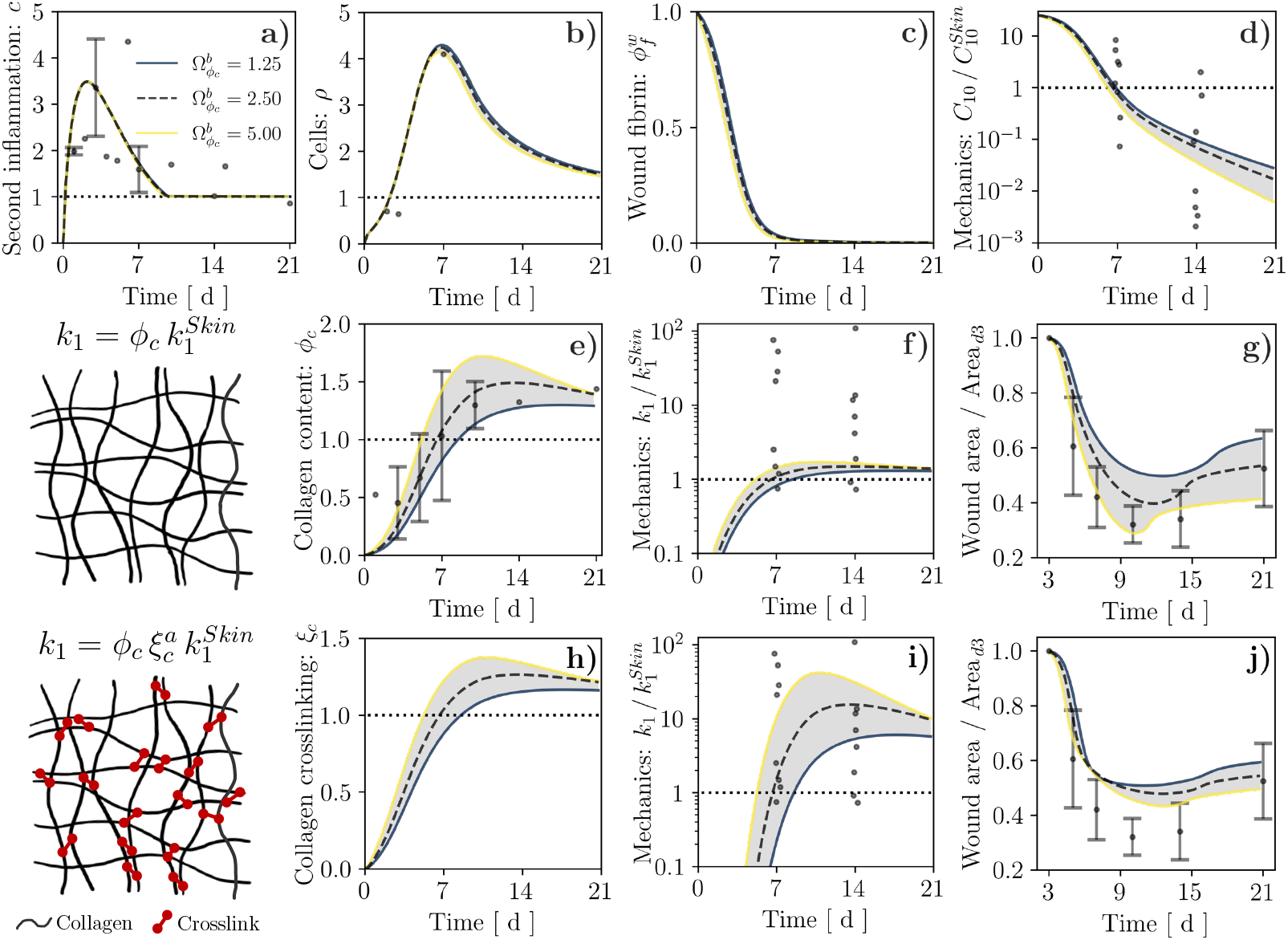
Results of wound healing simulation over a 21-day period for alternative links between the tissue collagen content, *ϕ*_*c*_, and the mechanical parameter *k*_1_, and alternative values of the parameter 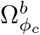 that controls collagen production by cells. (**a**–**d**) Temporal evolution of second inflammatory signal, *c*, cell population, *ρ*, wound fibrin content, 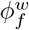, and mechanical parameter 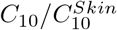 for either considered constitutive link. These quantities do not depend explicitly on *ϕ*_*c*_, hence we report them only once. (**e**–**g**) Temporal evolution of tissue collagen content, *ϕ*_*c*_, mechanical parameter 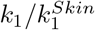, and wound area change resulting from assuming that *k*_1_ is proportional to *ϕ*_*c*_. (**h**–**j**) Temporal evolution of collagen crosslinking, *ξ*_*c*_, mechanical parameter 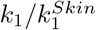, and wound area change resulting from assuming that *k*_1_ depends nonlinearly on *ϕ*_*c*_ via *ξ*_*c*_. Dots and error bars in (**a**,**b**,**e**,**g**,**j**): mean ± standard deviation of previously-published experimental data, *cf*. S1 Appendix and Ref. [10]. Dots in (**d**,**f**,**i**): values of 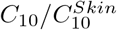 and 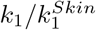 obtained from Bayesian calibration, *cf*. Fig. 3.

In Eq. (15), 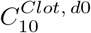 is the value of *C*_10_ in the wound right after the blood clot has formed, such that 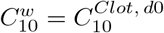 when 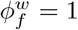 and 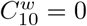 when 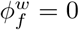. Conversely, *C*_10_ in the surrounding unwounded tissue is assumed constant throughout healing and set equal to the baseline skin value 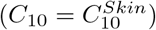. On the other hand, 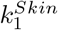 is the value of *k*_1_ for an unwounded tissue, such that 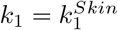 when *ϕ*_*c*_ = 1 (physiological value).

To address the plausibility of the constitutive hypotheses in Eqs. (14, 15), we simulate the evolution of *C*_10_ and *k*_1_ throughout wound healing (*cf*. S2 Video), in relation to the respective baseline values, 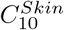 and 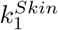. Notably, selecting 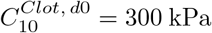, which is in line with previously-reported shear modulus values for venous thrombi [71], allows capturing the experimentally-informed evolution of 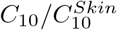 in the wound (Fig. 6d), supporting a dependence on 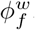. Conversely, the strong increase in *k*_1_ throughout healing, which we inferred from Bayesian parameter calibration, is not adequately captured by a proportional dependence on *ϕ*_*c*_. Indeed, even when varying the parameter 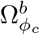 in the range of 0.5–2.0× to account for possible variability in collagen production (Fig. 6e), the predicted values of *k*_1_ hardly exceed the unwounded baseline (Fig. 6f). Importantly, this limitation does not affect the model ability to represent the temporal evolution of the diffusible fields *c* (Fig. 6a) and *ρ* (Fig. 6b), or its ability to capture visible wound area changes (Fig. 6g).

Based on the inadequacy of a proportional link between *k*_1_ and *ϕ*_*c*_, we hypothesize that this constitutive relation should additionally account for the progressive maturation of the newly-formed collagen network. Indeed, alterations in the degree and type of crosslinking have been reported to affect the emergent mechanical behavior of soft tissues such as tendons [72], uterine cervix [73], and skin wounds [10]. For simplicity, we focus on the degree of network crosslinking and consider a nonlinear relation between *k*_1_ and *ϕ*_*c*_, mediated by a crosslinking agent, *ξ*_*c*_:

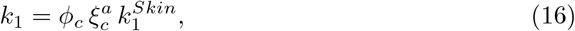

where *a* ≥ 1 is a phenomenological exponent controlling the degree of tissue nonlinearity associated with a given crosslinking. To prescribe the temporal evolution of *ξ*_*c*_, which constitutes an additional microstructural field in our model, we reason that crosslink formation should be positively correlated with *ϕ*_*c*_, since a higher collagen content should provide increased opportunities for physico-chemical interactions, and that the degree of network crosslinking cannot increase indefinitely with *ϕ*_*c*_ but should exhibit a saturating behavior given the finite size of crosslinks. On the other hand, the amount of crosslinks could be reduced spontaneously, as a consequence of stochastic unbinding [74], or indirectly, via depletion of some fibers within the ECM [75]. Accordingly, we express the source term for *ξ*_*c*_ as:

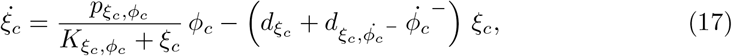

where 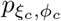 defines the natural forward rate of collagen crosslinking, 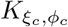 determines the saturation of *ξ*_*c*_ in reponse to 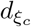 is the spontaneous decay rate for crosslinks, selected to ensure homeostasis of *ξ*_*c*_ in unwounded physiological conditions, and 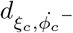 defines the relation between collagen fiber depletion and the associated crosslink depletion, which relates to the average number of crosslinks per collagen fiber.

Under the constitutive hypotheses in Eqs. (16, 17), collagen crosslinking increases concomitantly with *ϕ*_*c*_ during wound healing, from its initial value of 0 at the onset of wound healing to a supra-physiological value of 1.1–1.4× (depending on the value of 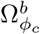) that is mostly preserved throughout days 10–21 post-wounding (Fig. 6h). Selecting a value *a* = 10 for the exponent in Eq. (16) allows our model to capture the strong increase and subsequent stabilization of *k*_1_ during the proliferation and remodeling stages of healing (Fig. 6i), while continuing to recapitulate previously-measured changes in the visible wound area (Fig. 6j). Importantly, the temporal evolution of the diffusible biochemical fields, of the fibrin content, and of the mechanical parameter *C*_10_ in the wound are largely unaffected by the assumed link between *k*_1_ and *ϕ*_*c*_ (*cf*. S3 Video), since the only possible dependence of these quantities on *k*_1_ is through the mechanical deformation of the wounded tissue, which we have already established to have a minor influence on *ρ* and *ϕ*_*c*_ in our model. Also note that the nonlinear relation between *k*_1_ and *ϕ*_*c*_ results in an increased sensitivity of *k*_1_ to the modeling parameter 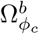 (Fig. 6i *vs*. Fig. 6f), allowing us to ascribe at least part of the experimental variability in *k*_1_ to potential specimen-specific differences in terms of collagen content and its degree of crosslinking.

### A closer look at mechano-biological signals: stretch *vs*. stiffness

Thus far, we have discussed the influence of wound deformability on the healing outcome and linked the emergent mechanical parameters to the biochemical and microstructural fields. In both cases, we have assumed that cell proliferation and collagen deposition depend on the elastic tissue deformation, as measured by *θ*^*e*^, via the function *Ĥ*. Selecting the parameters of *Ĥ* according to our previous works [37, 38] and to match available experimental data on the influence of stretching on fibroblast proliferation [66] yielded a surprisingly modest sensitivity of *ρ* and *ϕ*_*c*_ on *θ*^*e*^, *cf*. Fig. 5e,f and Fig. 6b,c. Since fibroblasts are known to be mechanosensitive, we now turn our attention to the mechano-biological pathway that links cell function to the mechanical state of the ECM. Specifically, we progressively increase the strength of this coupling to test its effect. Note that, for each considered value of Ω^*m*^, we also adjust the coefficients *c*_*tgt*_, *d*_*ρ*_, and *ϕ*_*c*_ so that physiological homeostasis is achieved in the unwounded tissue.

Surprisingly, increasing the value of 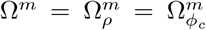 in Eqs. (6,7) appears to strongly mitigate fibrosis, leading to a marked decrease in *ρ, ϕ*_*c*_, and *ξ*_*c*_ (Fig. 7b,d,e). In turn, reduced cell infiltration mitigates the active stresses and the corresponding elastic pre-stretch in the wound (*θ*^*e*^, *cf*. Fig. 7c), leading to decreased area reduction over time (Fig. 7f). Slower cell infiltration in the wound also delays fibrin degradation, resulting in a slower decay for *C*_10_ (Fig. 7g). Concomitantly, the reduction in *ϕ*_*c*_ and *ξ*_*c*_ determines a marked decrease in *k*_1_ (Fig. 7h), which does not even reach the unwounded baseline value when the mechano-biological feedback is increased to Ω^*m*^ = 0.8, *cf*. S4 Video. Moreover, the emergent mechanical behavior of mature scar tissue (day 21 post-wounding) under uniaxial tensile conditions, evaluated analytically using Eq. (2), appears significantly softer with larger Ω^*m*^ (Fig. 7i). Taken together, these results indicate that positing a primary dependence of cell activity on ECM deformation might not allow capturing the onset of scar fibrosis. Importantly, this result follows from imposing an initially stress-free fibrin clot, implying an initially much smaller elastic deformation in the wound compared to the physiological state (Fig. 7c) and yielding sub-physiological values for *Ĥ*. As shown in S2 Appendix, this leads the source term *s*_*ρ*_ to decrease when Ω^*m*^ is increased if *ρ* ≥ 1, as also reflected by the trends in Fig. 7.

**Figure 7.**
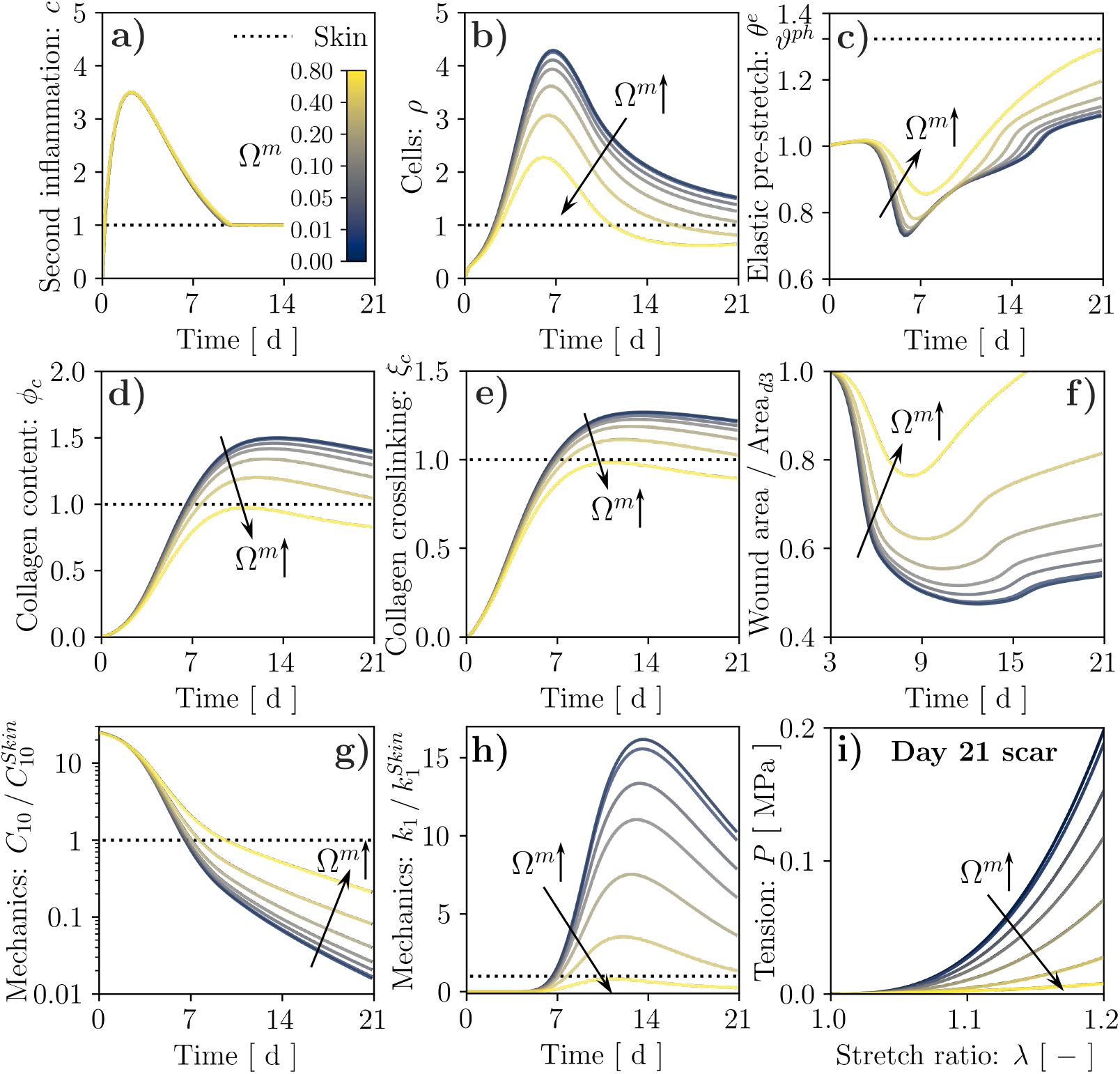
Results of wound healing simulation over a 21-day period for stretch-mediated mechanosensitivity and alternative values of the coupling strength, as controlled by the parameter Ω^*m*^. Temporal evolution of: second inflammatory signal, *c*, (**a**); cell population, *ρ*, (**b**); tissue elastic stretch, *θ*^*e*^ (**c**); tissue collagen content, *ϕ*_*c*_, (**d**); collagen crosslinking, *ξ*_*c*_, (**e**); wound area change (**f**); mechanical parameters 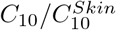 (**g**) and 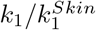 (**h**). The wound healing outcome in terms of tissue mechanical behavior is visualized by evaluating its tensile response at day 21 post-wounding (**i**).

To capture the enhanced tissue fibrosis that would be expected when strengthening the mechano-biological coupling [23, 24], we revisit the definition of *Ĥ* according to the widely accepted notion that stiffness can play a major role in regulating fibroblast activity [76–78]. Specifically, we assume that the mechanical parameter *k*_1_ provides a proxy of tissue stiffness and adopt once again the same definition for *Ĥ*_*ρ*_ and 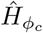 :

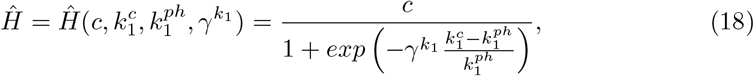

where 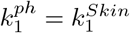 is the physiological value of *k*_1_, *i*.*e*. that of unwounded skin, 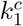 is the local value of *k*_1_, influenced by collagen deposition and crosslinking (Eq. (16)), and 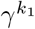 controls the slope of *Ĥ* around its midpoint, 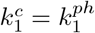, with a role analogous to that of *γ*^*e*^ in Eq. (9). Unlike Eq. (9), Eq. (18) also depends on *c*, linking the mechanosensitivity of cell proliferation and collagen deposition to ECM inflammation and reflecting the involvement of inflammatory pathways in tissue fibrosis [23].

As visible in Fig. 8, the mechanobiological coupling encoded by Eq. (18) induces a marked increase in *ρ, ϕ*_*c*_, and *ξ*_*c*_ (Fig. 8b,d,e and S5 Video) when the value of 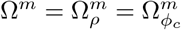 is increased, contrary to what observed in Fig. 7. The higher cell density also yields larger active stresses and stronger ECM contraction (Fig. 8c), enhancing the wound area reduction over time (Fig. 8f and S5 Video) and causing faster depletion of the fibrin clot, which results in a more rapid decay of *C*_10_ (Fig. 8g and S5 Video). Concomitantly, the increase in *ϕ*_*c*_ and *ξ*_*c*_ determines a marked increase in *k*_1_ (Fig. 8h and S5 Video) and a corresponding stiffening of the emergent mechanical behavior for a 21-day-old scar tissue (Fig. 8i and S5 Video). Unlike for the previous mechanobiological coupling, the *k*_1_ increase resulting from a larger value of Ω^*m*^ now triggers a positive feedback loop encoded by Eq. (18). In fact, for the extreme case of Ω^*m*^ = 0.8 (S6 Video), this loop causes *ρ, ϕ*_*c*_, *ξ*_*c*_, and thus *k*_1_, to maintain sustained overexpression with respect to their physiological baseline values. This aspect is further analyzed in S3 Appendix, where we examine the equilibrium points of an ODE system derived from Eqs. (6, 7, 16–18), which can be considered representative of a 0-dimensional tissue region without biochemical field diffusion (Eq. (3)) or remodeling (Eq. (8)). Our analysis shows that Ω^*m*^ affects the number of equilibrium points for the system, and setting Ω^*m*^ = 0.8 leads to a bi-stable system. Thus, the wound can reach a supra-physiological steady state, indicative of permanent fibrosis, while the surrounding unwounded tissue evolves toward the physiological steady state.

**Figure 8.**
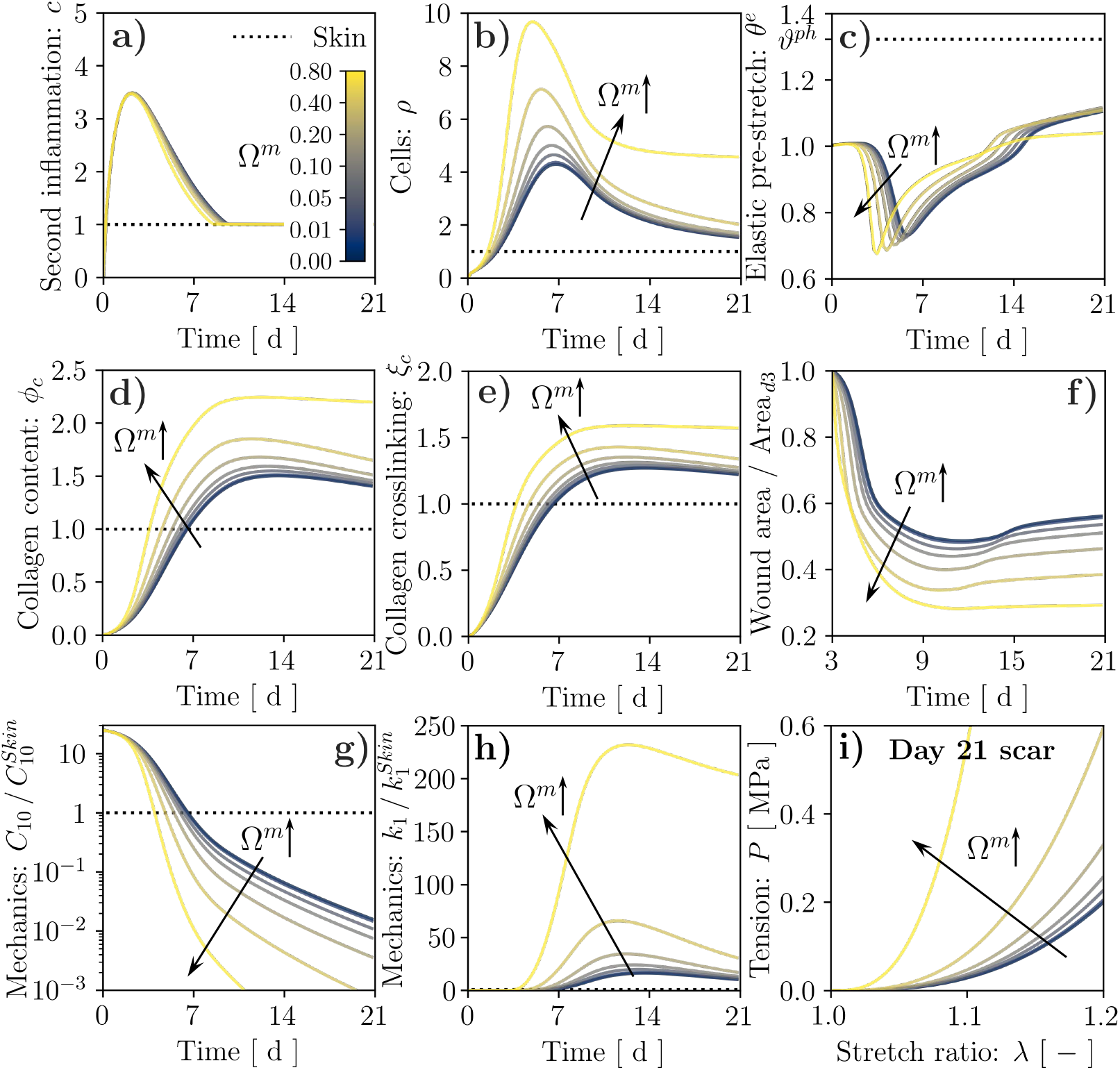
Results of wound healing simulation over a 21-day period for stiffness-mediated mechanosensitivity and alternative values of the coupling strength, as controlled by the parameter Ω^*m*^. Temporal evolution of: second inflammatory signal, *c*, (**a**); cell population, *ρ*, (**b**); tissue elastic stretch, *θ*^*e*^ (**c**); tissue collagen content, *ϕ*_*c*_, (**d**); collagen crosslinking, *ξ*_*c*_, (**e**); wound area change (**f**); mechanical parameters 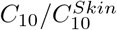 (**g**) and 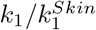 (**h**). The wound healing outcome in terms of tissue mechanical behavior is visualized by evaluating its tensile response at day 21 post-wounding (**i**).

## Discussion

Computational models of cutaneous wound healing are gaining increasing popularity as promising tools in bioengineering and clinical contexts, *e*.*g*. personalized medicine [29,79] and *in silico* clinical trials [80]. However, recent advances in the representation of the biochemical processes underlying tissue repair have not been paralleled by similar developments in the description of the wound mechanics, mainly hindered by scant experimental evidence. In this study, we proposed to overcome these limitations by leveraging one of the very few available experimental datasets on the evolution of murine wound biomechanics throughout healing [39].

In order to determine constitutive model parameters for wounded and unwounded skin, and quantify their variability, we have established a novel hierarchical Bayesian inverse analysis procedure that is broadly applicable towards determining sets of inter-related, specimen-specific mechanical parameters from corresponding experimental data (Figs. 2–3). Despite the large variability intrinsic to biological tissue properties, our approach allowed identifying overall trends for the wound constitutive parameters, highlighting clear softening of the non-collagenous ground substance throughout healing (Fig. 3a) and sustained stiffening of the collagenous ECM with respect to the unwounded baseline (Fig. 3b); we interpret the latter as indicative of wound/scar fibrosis.

Aiming to establish a versatile *in silico* tool to test alternative hypotheses on the bio-mechanical and mechano-biological pathways involved in wound healing, we then calibrated our systems bio-chemo-mechanobiological FE model [37,40] to recapitulate the temporal evolution of several key biochemical and morphological aspects of murine wound healing over a 21-day period (Fig. 5). Altering the tissue biomechanical parameters according to the 95% CI obtained from the Bayesian calibration procedure allowed us to assess their influence on the healing outcome in terms of wound permanent contracture and visible area changes. Specifically, we observed that softer/stiffer wounds develop into smaller/larger scars with reduced/increased permanent deformation (Fig. 5), highlighting a cell-mediated mechanism whereby the wound ECM mechanics influences the outcome of the tissue repair processes.

Next, we used our model to propose bio-mechanical constitutive links for the emergent mechanical parameters of wounded tissue (*C*_10_, *k*_1_), starting from the underlying microstructural protein content 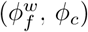, whose spatial and temporal evolution is determined by the biochemical (*c, ρ*) and mechanical (*θ*^*e*^, *θ*^*p*^) fields. In stark contrast with several previous works [67–70], we found that a simple proportional relation between tissue stiffness and collagen content is unable to explain the changes in the mechanical parameters *k*_1_ that we derived from experimental data. Conversely, a nonlinear relation captures the strong increase and subsequent stabilization of *k*_1_ during the proliferation and remodeling stages of healing (Fig. 6), and allows ascribing at least part of the experimental variability in *k*_1_ to potential specimen-specific differences in *ϕ*_*c*_ and *ξ*_*c*_. In line with previous works on the mechanics of collagen and actin networks, whose emergent response depends nonlinearly on crosslink density [81–83], our constitutive hypothesis was based on a power law, which we speculate to have a microstructural origin. Indeed, murine wound/scar tissue has been reported to feature an increased proportion of collagen crosslinks associated with fibrotic tissue with respect to the unwounded baseline [10, 84]. Similarly, the ratio between such crosslinks and those typical of soft connective tissues was upregulated in idiopathic pulmonary fibrosis, as were the density of mature crosslinks and the tissue stiffness, but not the collagen content [85]. Further supporting our modeling approach, the mature/immature crosslink ratio correlated positively with changes in the mechanical stiffness of lateral collateral ligament following injury [72], suggesting a dominant role of tissue ‘quality’ over its ‘quantity’ towards determining tissue fibrosis.

Lastly, we have leveraged our model to analyze the mechano-biological coupling from the mechanical state of the wound to the fibroblast proliferation and collagen deposition. In line with ubiquitous clinical evidence of the role of mechanical forces within tissue fibrosis [23–26], we expected that increasing this coupling would enhance cell infiltration, collagen deposition, and thus increase *k*_1_ in the wound bed, ultimately exacerbating scar fibrosis. However, we found that a deformation-driven link between cell activity and the ECM mechanical state — supported by several works on stretch mechanosensitivity in fibroblasts [20, 21] — led to mitigated scarring, owing to the sub-physiological deformations of the wound ECM (Fig. 7). On the other hand, extensive recent work in cell mechanobiology has highlighted a strong sensitivity of fibroblast activity to ECM stiffness [76–78, 86] through a pathway involving integrin-mediated adhesion [77, 87, 88], which we have also recently used to explain stretch-mediated mechanosensitivity in cells by combining Bell’s adhesion kinetics with the typical nonlinear strain-stiffening of collagenous ECMs [89]. Indeed, considering stiffness-driven cell mechanosensitivity (Fig. 8) led all markers of tissue fibrosis included in our model (*ϕ*_*c*_, *ξ*_*c*_, *k*_1_) to be overexpressed when increasing the coupling strength, suggesting that the nature of fibroblast mechanosensing and its involvement in wound healing remain open questions. Interestingly, for the strongest coupling that we considered (Ω^*m*^ = 0.8), the wound and the surrounding skin evolved towards different steady states, suggesting irreversible changes in the scar tissue. This result, obtained by analyzing bifurcations in the ODE system comprising the key evolution equations of our FE model (S3 Appendix), matches the evidence that injured tissues can never regain the properties of native skin [10, 13–15]. This study explored several *what if* scenarios that challenge our fundamental understanding of the interplay between the biological, chemical and physical events involved in wound healing. However, it is not without limitations. First, while we have informed the model with experimental data, we have also neglected a few key aspects of wound healing. We focused on the evolving mechanics of the rebuilding dermal tissue, ignoring the role of keratinocytes within wound epithelialization and that of endothelial cells within angiogenesis. Both cell types are known to strongly affect the outcome of wound healing, *e*.*g*. by stimulating fibroblast function [18,90], so that including them in our model would contribute to a broader and deeper understanding of the tissue repair process. Second, we have ignored any possible role of tissue anisotropy and three-dimensional geometries — both in the wound and in the surrounding skin — due to the lack of corresponding experimental information. Future experimental investigations of wound healing biomechanics should specifically address these aspects, providing invaluable quantitative data for further model developments. We have also considered a continuum representation of the tissues, which can provide an accurate description of their mechanics but is only one of the strategies for modeling wound healing. Since biological processes such as cell mechanosensing might be better captured by computational models at smaller scales, *e*.*g*. discrete fiber network and agent based models, a natural future development of this work is to include a coupled multi-scale approach. Finally, a key result of this study is the proposition of a nonlinear link between the emergent mechanical response of the wound tissue and its collagen content. While this is not surprising, given the known network-like characteristics of collagenous tissues, the exponent *a* that we have used in Eq. (16) was selected arbitrarily and with the specific goal of matching the available experimental data. Future work should focus on including a physically-based link between the emergent mechanical behavior of the newly-formed collagenous networks and microstructural parameters such as the ratio between various collagen isoforms, the density, type, and kinetics of crosslinks, relevant network statistics (*e*.*g*. fiber diameters, stiffness, length between crosslinks), as well as the possible mechanical role of non-collagenous proteins (*e*.*g*. proteoglycans and glycoproteins).

## Conclusion

Motivated by its potential relevance within bioengineering and clinical contexts, we have presented a calibrated systems bio-chemo-mechanobiological FE model of wound healing progression that accounts, for the first time, for the local changes in stiffness of the wounded tissue. The time-evolving mechanical characteristics of the repairing skin were inferred based on a novel, broadly applicable Bayesian inverse analysis procedure. The uncertainty propagation step following calibration allowed us to investigate the direct dependence between the local changes in wound stiffness to macroscopic contraction upon healing. The versatility of our model towards formulating and testing biomechanical and mechanobiological hypotheses was demonstrated by evaluating alternative links between the wound microstructural composition and its emergent mechanical behavior, as well as by discussing the implications of stretch- *vs*. stiffness-dominated mechanobiological coupling towards explaining the onset of irreversible scar fibrosis.

## Supporting information

**S1 Fig.**
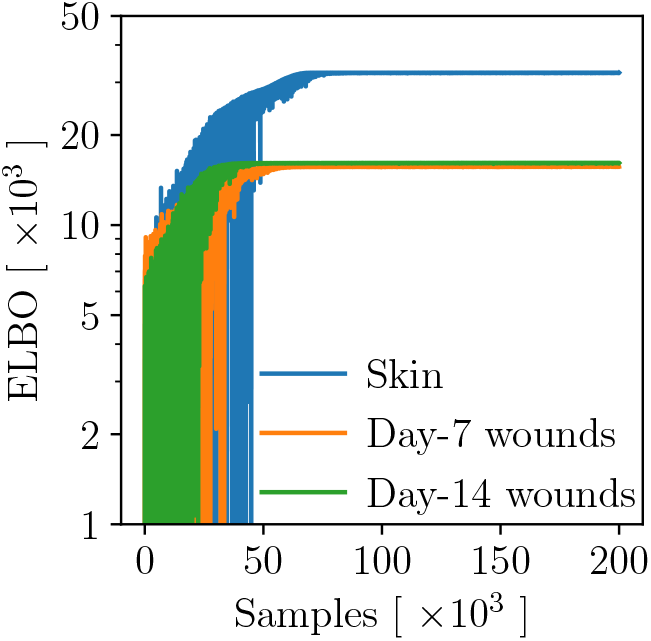
Convergence of ELBO for hierarchical Bayesian model calibrations.

**S2 Fig.**
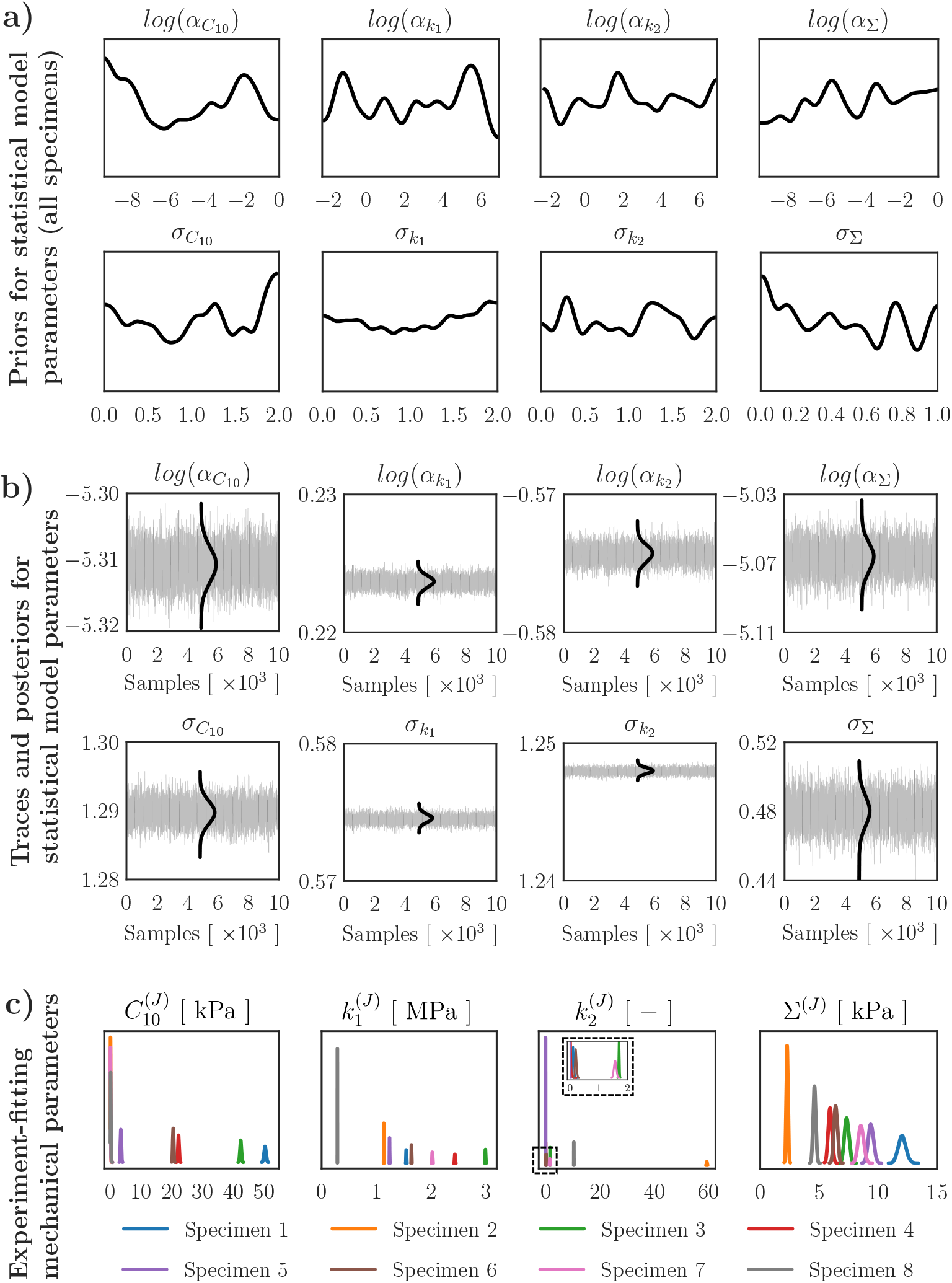
Hierarchical Bayesian model calibration for unwounded skin.

**S3 Fig.**
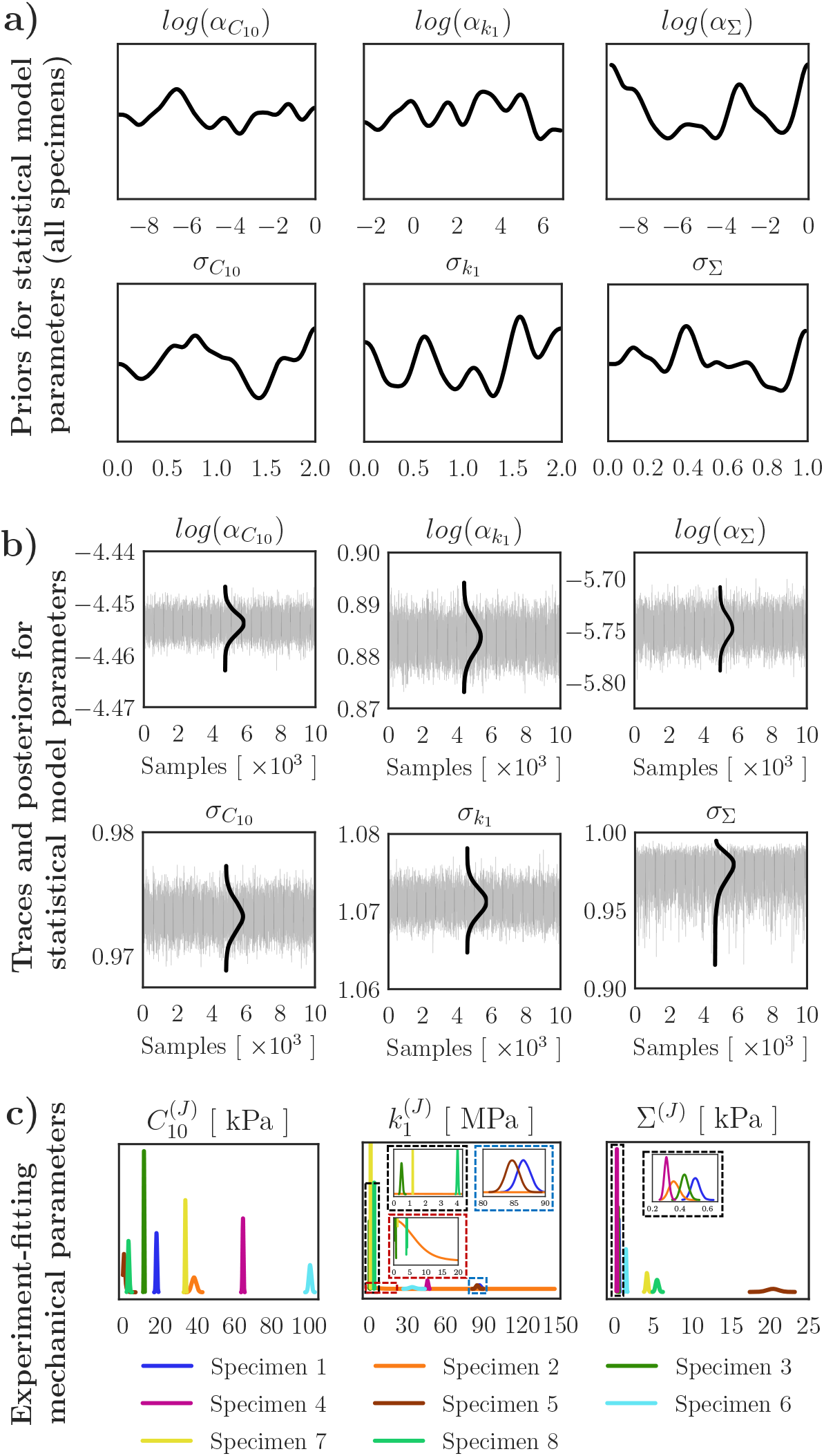
Hierarchical Bayesian model calibration for 7-day-old wounds.

**S4 Fig.**
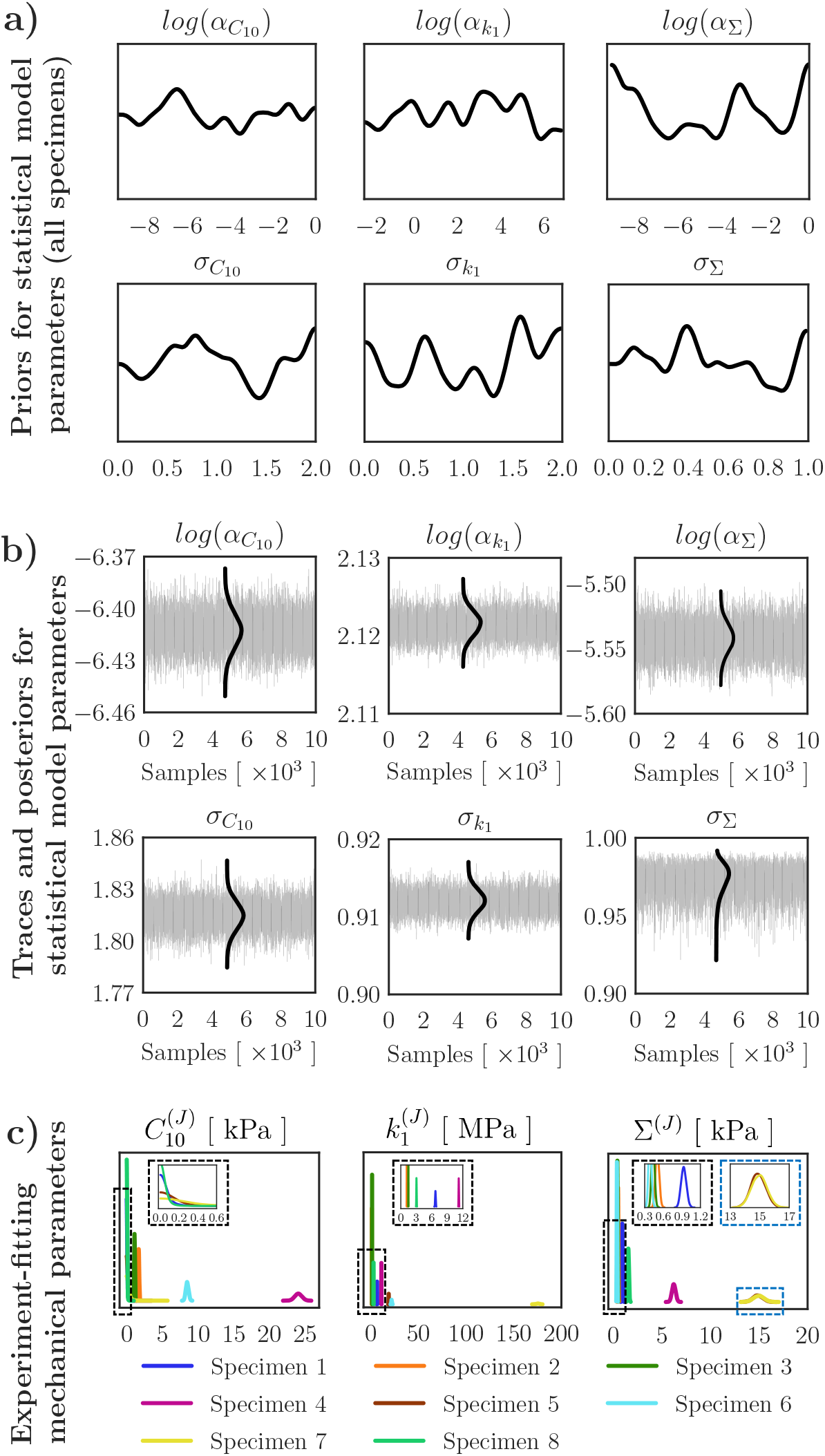
Hierarchical Bayesian model calibration for 14-day-old wounds.

**S1 Video. Simulation of wound healing progression with hard-coded mechanical parameters evolving linearly between the experimentally-informed median values, and strain-driven mechanosensing with Ω**^***m***^ **= 0.01**.

**S2 Video. Simulation of wound healing progression assuming that the mechanical parameter** *k*_1_ **depends linearly on the collagen content, and strain-driven mechanosensing with Ω**^***m***^ **= 0.01**.

**S3 Video. Simulation of wound healing progression assuming a power law linking the mechanical parameter *k***_**1**_ **to the collagen content, and strain-driven mechanosensing with Ω**^***m***^ **= 0.01**.

**S4 Video. Simulation of wound healing progression assuming a power law linking the mechanical parameter *k***_**1**_ **to the collagen content, and strain-driven mechanosensing with Ω**^***m***^ **= 0.8**.

**S5 Video. Simulation of wound healing progression assuming a power law linking the mechanical parameter *k***_**1**_ **to the collagen content, and stiffness-driven mechanosensing and Ω**^***m***^ **= 0.01**.

**S6 Video. Simulation of wound healing progression assuming a power law linking the mechanical parameter *k***_**1**_ **to the collagen content, and stiffness-driven mechanosensing and Ω**^***m***^ **= 0.8**.

## S1 Appendix. Literature data for wound biochemistry and mechanobiology

To perform wide-range calibration of our custom FE model, we derive temporal evolutions of the biochemical fields *c, ρ*, and *ϕ*_*c*_ from published data on full-thickness excisional wounds in wild type mice (diameter: 3–8 mm). Unless numerical values are explicitly reported, we extract them from published charts using the web-based tool ‘WebPlotDigitizer’ [91]. For studies featuring measurement replicates, we focus on their average at each available time point. When multiple data from different studies are available for the same time point, we quantify and display their average and standard deviation (*cf*. Figs. 5-6 and Fig. A1.1).

To inform the temporal evolution of *c*, we consider TGF-β1 as a reliable indicator of the second inflammatory wave progression, owing to its undisputed role as the growth factor with the broadest spectrum of actions on cell activity within wound healing [59]. Focusing on experimental studies reporting the evolution of TGF-β1 throughout healing and the corresponding baseline values in unwounded tissue [57–60], we obtain the temporal evolutions in Fig. A1.1a.

For the cell content, *ρ*, we focus on previous studies reporting the number of fibroblasts in wounded and unwounded tissue [57, 61], and obtain the temporal evolutions in Fig. A1.1b.

For the collagen content, *ϕ*_*c*_, we consider measurements of wound hydroxyproline content [62–65] and obtain the corresponding collagen amount per mass of wet tissue according to the relation:

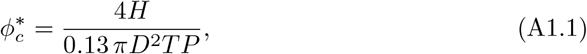

where *H* is the hydroxyproline mass measured in a tissue biopsy of diameter *D*, thickness *T*, and density *P*, while 0.13 is a conversion factor corresponding to the typical percentage of hydroxyproline within collagen [92]; in line with previous measurements, we take *P* = 1.1 mg mm^−3^ [93]. Using a similar approach, we also estimate the average collagen content in unwounded murine skin to be 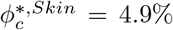 [64, 94–99], leading to the normalized wound collagen contents shown in Fig. A1.1c and in Figs. 5-6.

For the dependence of cell activity on tissue deformation, we consider previous *in vitro* data on the proliferation of human patellar tendon fibroblasts under uniaxial cyclic stretch of increasing magnitude [66]. For comparability with our study, where we simulate tissue biaxial stretching from a physiological deformation state, we assume that the data in Ref. [66] can be interpreted as representative of fibroblast overproduction induced by a tissue overstretching *θ*^*e*^*/θ*^*ph*^ ∼ (*λ*^*over*^*/λ*^*ref*^)^2^, where *λ*^*over*^ are the uniaxial stretch values used in Ref. [66] and *λ*^*ref*^ = 1 is the corresponding reference value (no stretch). To determine the overexpression of *ρ* for an unwounded tissue under stretch, we consider the ODE system comprising Eqs. (5–7) and predict the values of *c, ρ*, and *ϕ*_*c*_ after 210 days of application of an areal deformation *θ*^*e*^. Specifically, we set *α* = 0 and select all parameters except for 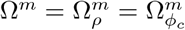 according to S3 Table and S4 Table. Since we are interested in stretch-mediated mechanosensing, we adopt the definition of *Ĥ* in Eq. (9). As shown in Fig. A1.2, increasing/decreasing the tissue stretch around its physiological value results in increased/decreased fibroblast production, in a way that depends on the value of Ω^*m*^. Since values of Ω^*m*^ in the range of 0.005 − 0.02 well capture the experimental data in Ref. [66], we select Ω^*m*^ = 0.01 as the reference value for this study.

**Figure A1.1.**
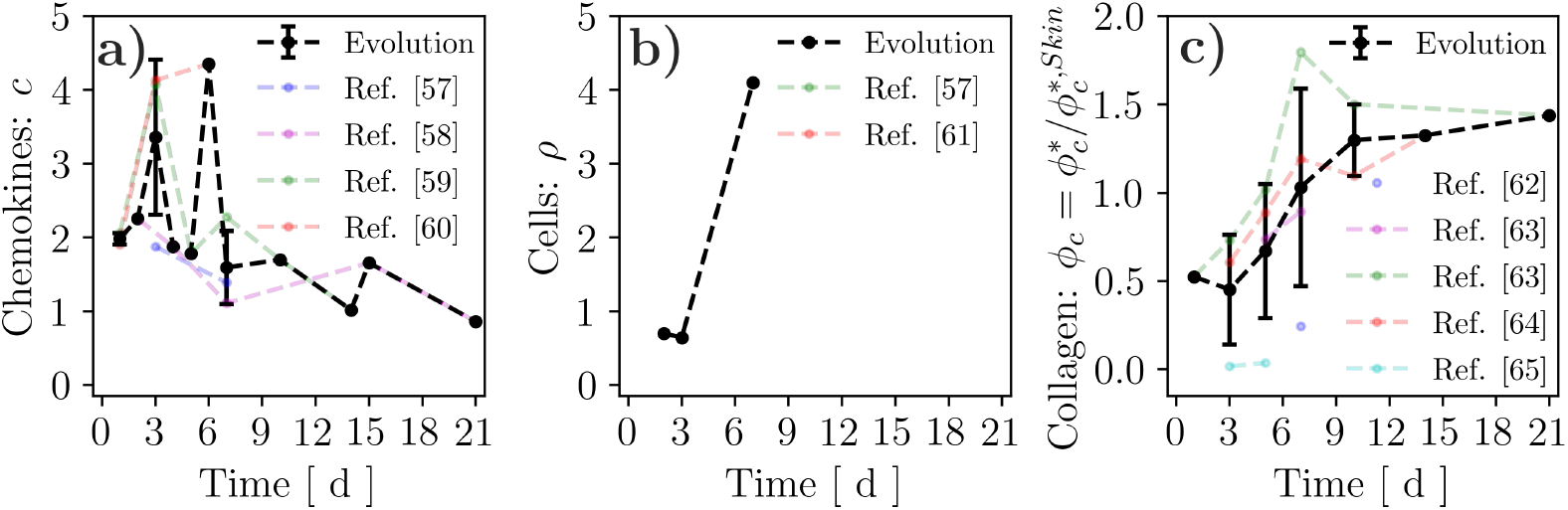
Temporal evolution of cytokines, *c*, cells, *ρ*, and collagen content, *ϕ*_*c*_, according to several published studies (colored translucent dots connected by dashed lines showing trends). The data points for comparison with simulations are obtained by averaging information at corresponding time points obtained from different studies (black dots and error bars: mean ± standard deviation; dashed lines show trends).

**Figure A1.2.**
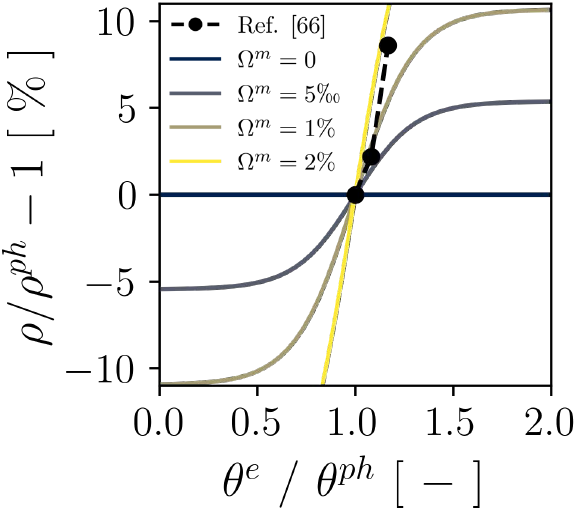
Dependence of fibroblast production on tissue stretch for alternative values of the coupling parameter Ω^*m*^, in comparison with experimental data (black dots connected by a dashed line).

## S2 Appendix. Sensitivity of *s*_*ρ*_ to Ω^*m*^ for sub-physiological elastic stretch

In Fig. 7, we have shown that assuming a stretch-mediated mechanobiological coupling yields a marked decrease in *ρ* and other variables in the wound when increasing the strength of the coupling (Ω^*m*^), a behavior that we have explained in terms of the sub-physiological elastic deformation in the wound that results from imposing an initially stress-free fibrin clot. Here, we further analyze the link between *ρ* and Ω^*m*^ by analytically computing the derivative of the source term in Eq. (6), *s*_*ρ*_, with respect to the parameter defining the coupling strength, Ω^*m*^:

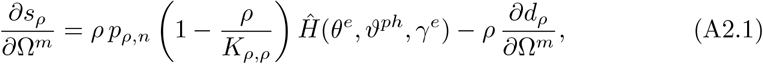

where *d*_*ρ*_ depends on Ω^*m*^ through the homeostasis constraint imposed considering an unwounded and physiologically-loaded tissue

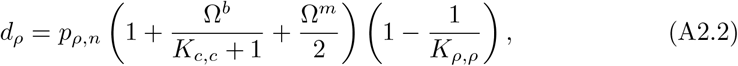

such that

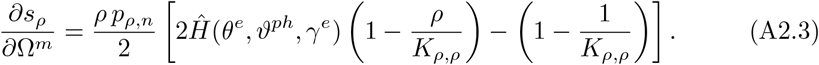

Since we are considering sub-physiological values of the elastic deformation, we can use the inequality *Ĥ* (*θ*^*e*^, *ϑ*^*ph*^, *γ*^*e*^) < 1*/*2 to write

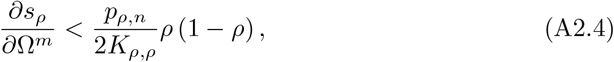

whose right-hand side term provides an upper bound for *∂s*_*ρ*_*/∂*Ω^*m*^ and has sign depending on *ρ*, indicating that *s*_*ρ*_ will decrease for increasing Ω^*m*^ for values of *ρ* ≥ 1 when *θ*^*e*^ < *ϑ*^*ph*^.

## S3 Appendix. Equilibrium points for ODE system comprising Eqs. (6,7,17)

In Fig. 8, we have observed that a sufficiently large value of Ω^*m*^, *e*.*g*. Ω^*m*^ = 0.8, induces sustained overexpression of *ρ, ϕ*_*c*_, *ξ*_*c*_, and *k*_1_ throughout the simulated wound healing time span. To provide an explanation for this peculiar behavior, we study the equilibrium of the ODE system comprising Eqs. (6,7,17), where the definitions of *k*_1_ and *Ĥ* are given in Eq. (16) and Eq. (18), respectively. This system of equations can be regarded as descriptive of a 0-dimensional tissue region lacking any biochemical field diffusion (Eq. (3)) or ECM remodeling (Eq. (8)). Note that we exclude Eq. (5) from the ODE system, and thus set *c* = 1, corresponding to the assumption of a negligible contribution of the secondary inflammation wave on the long-term tissue behavior. Focusing on the region of space *ρ* × *ϕ*_*c*_ × *ξ*_*c*_ = [0; 8] × [0; 3] × [0; 3], we set the model parameters to the values indicated in S3 Table and S4 Table and identify the loci of points satisfying the conditions *s*_*ρ*_ = 0, 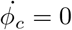, and 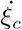, respectively. More explicitly, we solve each of the following nonlinear algebraic equations for alternative choices of Ω^*m*^, knowing that the system will be in equilibrium when all equations are satisfied:

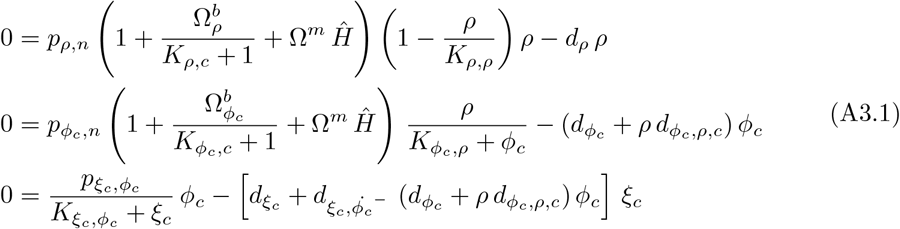

First, we consider the default value Ω^*m*^ = 0.01. As shown in Fig. A3.1a, the surfaces corresponding to *s*_*ρ*_ = 0 (blue), 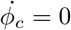 (green), 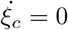 (red) intersect along multiple lines, but there are only 2 points where all of them are satisfied, *i*.*e*. where the system is in equilibrium. One is the trivial solution (*ρ, ϕ*_*c*_, *ξ*_*c*_) = (0, 0, 0), capturing the notion that a system without cells or collagen will not spontaneously accumulate any of these quantities nor crosslinks. Instead, the non-trivial solution (*ρ, ϕ*_*c*_, *ξ*_*c*_) = (1, 1, 1) corresponds to the physiological condition and will act as attractor for any tissue in supra-physiological conditions, indicating that the wound behavior shown in Fig. 8 for Ω^*m*^ = 0.01 will eventually evolve towards the physiological state of the unwounded tissue. Similar observations hold for the case Ω^*m*^ = 0.4, Fig. A3.1b. Conversely, when Ω^*m*^ = 0.8, the system exhibits 3 non-trivial and 1 trivial equilibrium points (Fig. A3.1c). This phase-space topology implies a bi-stable system, with one stable point continuing to be the physiological equilibrium, and the other stable point corresponding to a fibrotic state. To confirm this, we consider the results of our FE simulations for Ω^*m*^ = 0.8 and represent the streamlines on planes of constant *ξ*_*c*_, *ρ*, or *ϕ*_*c*_, both for the unwounded (far field) and wounded tissue at day 21 post-wounding. As visible in Fig. A3.1d–f, the equilibrium point at (1, 1, 1) acts as an attractor for the far field (black dot), which will thus eventually return to the physiological state. On the contrary (Fig. A3.1g–i), the day-21 values of *ρ, ϕ*_*c*_, and *ξ*_*c*_ place the wounded tissue (red dot) in a different landscape region, where the attractor is the point at (*ρ, ϕ*_*c*_, *ξ*_*c*_) = (6.1, 2.4, 1.7), leading the system towards a supra-physiological steady state that corresponds to permanent fibrosis.

**Figure A3.1.**
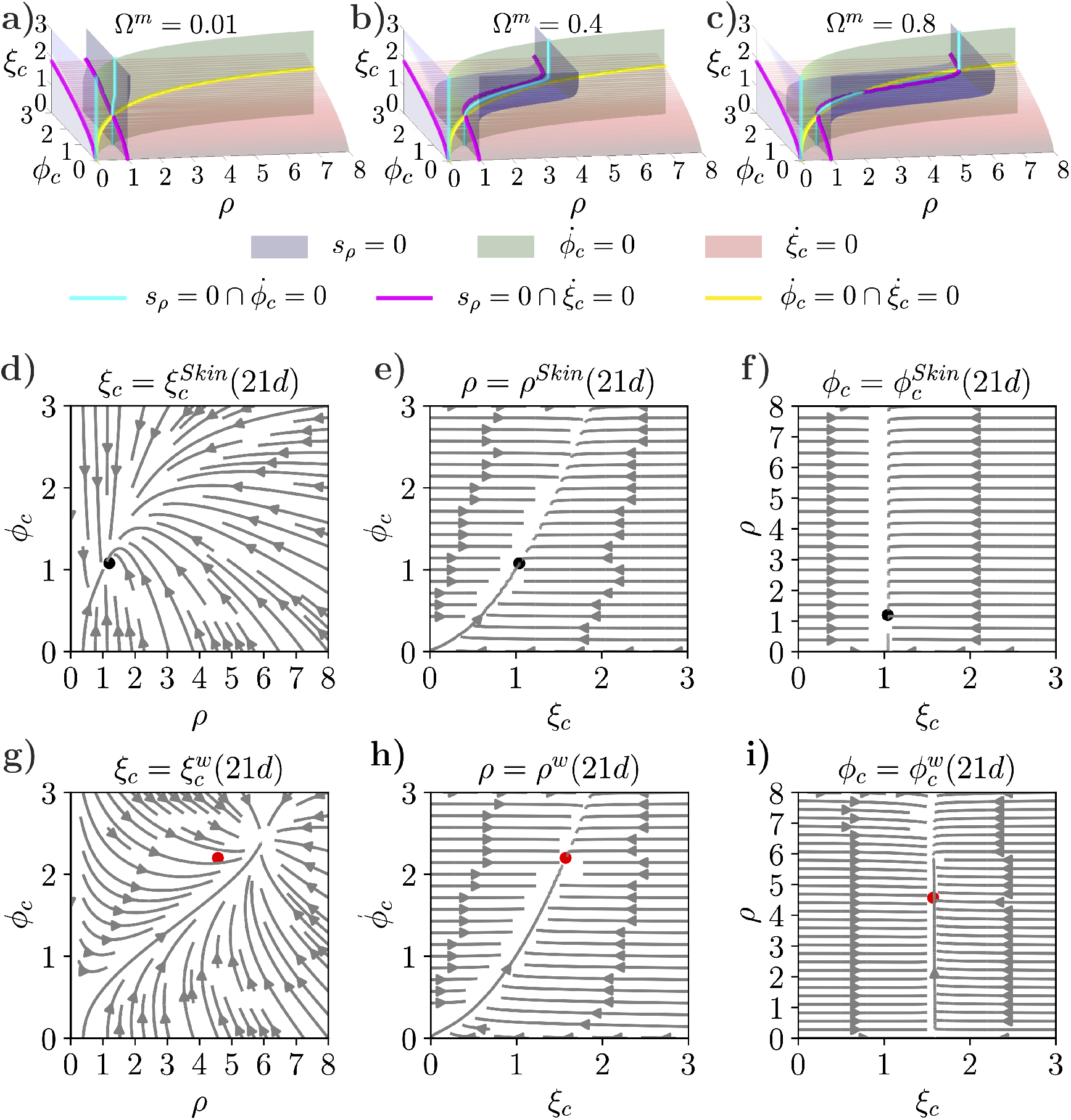
Stability analysis of an ODE system representative of long-term tissue progression. (**a**–**c**) Loci of points independently satisfying the conditions *s*_*ρ*_ = 0 (blue surface), 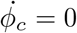 (green surface), and 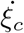 (red surface), as well as their intersections (cyan, magenta, and yellow lines). Line intersections identify equilibrium points for the ODE system. The phase-space topology, and thus the number of equilibrium points, is affected by the value of the coupling parameter Ω^*m*^. (**d**–**f**) Streamlines on planes of constant *ξ*_*c*_, *ρ*, or *ϕ*_*c*_ for unwounded skin tissue and Ω^*m*^ = 0.8. The black dot indicates the state of skin at day 21 post-wounding, as obtained from FE simulations. (**g**–**i**) Streamlines on planes of constant *ξ*_*c*_, *ρ*, or *ϕ*_*c*_ for wounded tissue and Ω^*m*^ = 0.8. The red dot indicates the state of the wound at day 21 post-wounding, as obtained from FE simulations.

---

**S1 Table.**
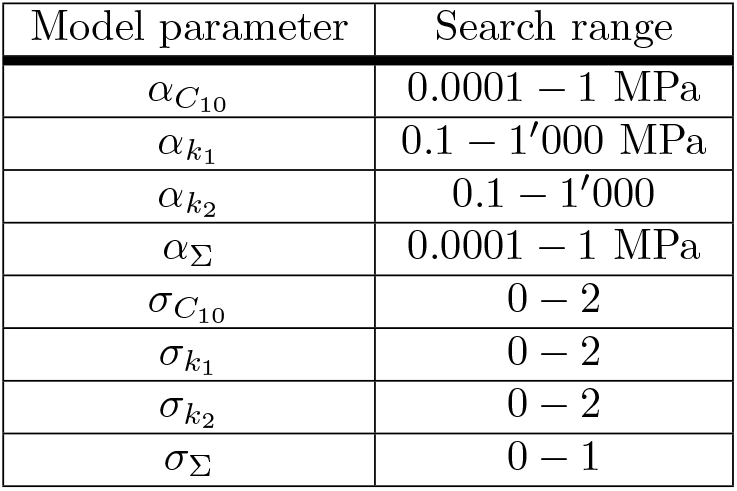
Search ranges for Bayesian model parameters.

| Model parameter | Search range |
| --- | --- |
| $\alpha_{C_{10}}$ | 0.0001 – 1 MPa |
| $\alpha_{k_1}$ | 0.1 – 1'000 MPa |
| $\alpha_{k_2}$ | 0.1 – 1'000 |
| $\alpha_{\Sigma}$ | 0.0001 – 1 MPa |
| $\sigma_{C_{10}}$ | 0 – 2 |
| $\sigma_{k_1}$ | 0 – 2 |
| $\sigma_{k_2}$ | 0 – 2 |
| $\sigma_{\Sigma}$ | 0 – 1 |

**S2 Table.** Global simulation parameters.

| Parameter | Value | Notes |
| --- | --- | --- |
| Total simulation time | 504 h = 21 d | From range of available data |
| Simulation time step | 0.05 h | Smaller than $\tau_{\lambda_a^p} = \tau_{\lambda_s^p}$ |
| Domain size in $X$: $L_X$ | 50 mm | $10D^w$ |
| Domain size in $Y$: $L_Y$ | 50 mm | $10D^w$ |
| Wound diameter: $D^w$ | 5.0 mm | Ref. [39] |
| Skin thickness | 1.7 mm | Ref. [10] |
| Physiological stretch along $x$: $\lambda_x$ | 1.15 | Within range from Refs. [51, 55, 56] |
| Physiological stretch along $y$: $\lambda_y$ | 1.15 | Within range from Refs. [51, 55, 56] |
| Physiological value of $\theta^e$: $\vartheta^{ph}$ | 1.3225 | $= \lambda_x \lambda_y$ |

**S3 Table.** Parameters for diffusible biochemical fields.

| Parameter | Value | Notes |
| --- | --- | --- |
| $D_{\alpha,\alpha}$ | 0.075 mm<sup>2</sup>/h | Assumed equal to $D_{c,c}$ |
| $d_{\alpha}$ | 0.01 1/h | Assumed equal to $d_c$ |
| $D_{c,c}$ | 0.075 mm<sup>2</sup>/h | Ref. [37] |
| $d_c$ | 0.01 1/h | Ref. [37] |
| $p_{c,\alpha}$ | 0.23 1/h | Selected to match peak in $c$ |
| $p_{c,\rho}$ | 1.5 1/h | Ref. [100] |
| $K_{c,c}$ | 0.5 | Assumed |
| $D_{\rho,\rho}$ | 0.035 mm<sup>2</sup>/h | Ref. [37] |
| $D_{\rho,c}$ | $8 \times 10^{-5}$ mm<sup>2</sup>/h | Ref. [37] |
| $p_{\rho,n}$ | 0.034 1/h | Ref. [37] |
| $\Omega_{\rho}^b$ | 5 | Selected to match peak in $\rho$ |
| $\Omega_{\rho}^m$ | 0.01 (Range: 0 – 0.8) | Varied in Figs. 7-8 |
| $\gamma^e$ | 5 | Ref. [37] |
| $\gamma^{k_1}$ | 0.016 | Assumed |
| $K_{\rho,\rho}$ | 30 | Selected to match peak in $\rho$ |
| $K_{\rho,c}$ | 10 | Selected to match peak in $\rho$ |
| $K_{f,c}$ | $10^{-5}$ | Ref. [37] |
| $f_{\rho,n}$ | 0.04 MPa/mm<sup>3</sup> | Ref. [37] |
| $\Omega_f^b$ | 1 | Ref. [37] |

**S4 Table.**
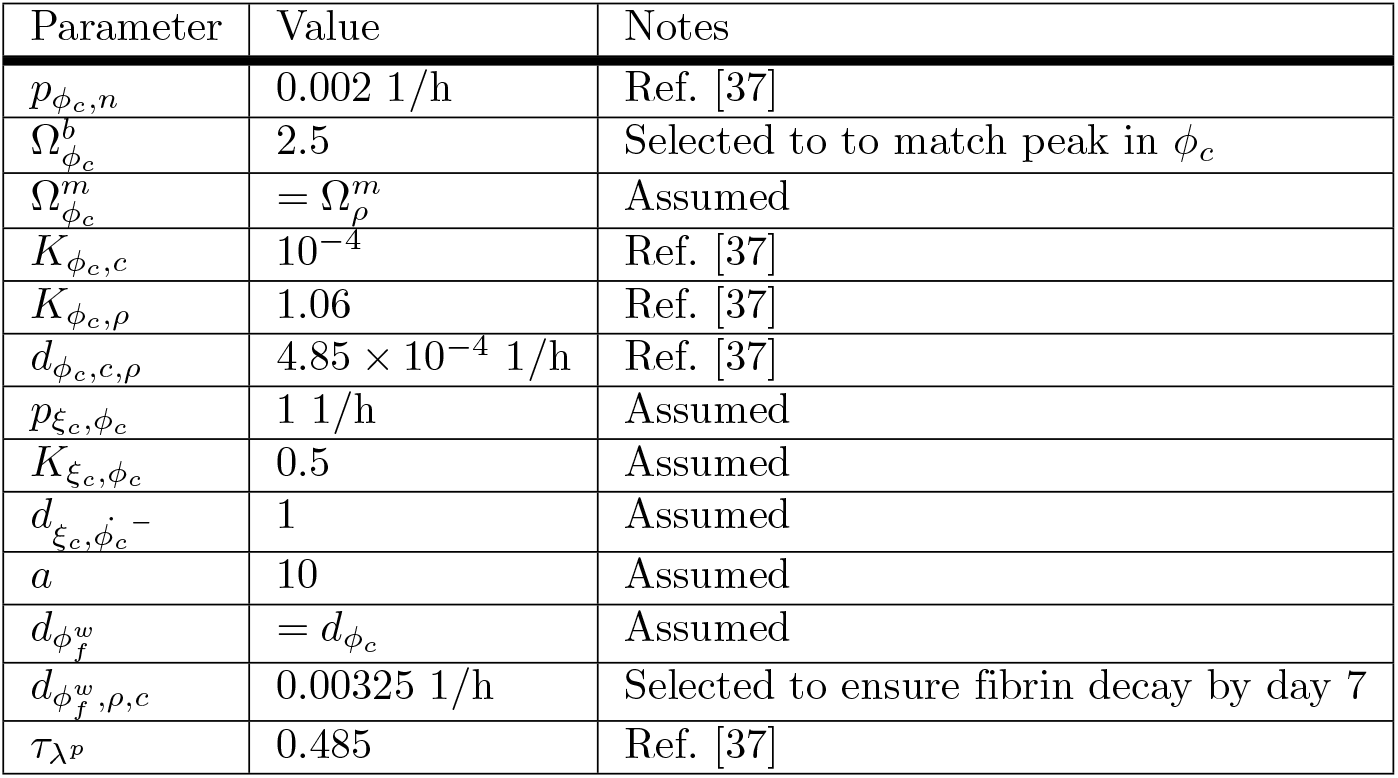
Parameters for microstructural fields.

**S5 Table.** Emergent constitutive biomechanical parameters.

| Parameter | Value | Notes |
| --- | --- | --- |
| $C_{10}^{Skin}$ | 12 kPa | Median value from Fig. 3a |
| $k_1^{Skin}$ | 1.6 MPa | Median value from Fig. 3b |
| $k_2^{Skin}$ | 0.88 MPa | Median value from Fig. 3c |
| $C_{10}^{i.w.}$ | 0.0 kPa | Due to tissue removal as part of wounding |
| $k_1^{i.w.}$ | 0.0 MPa | Due to tissue removal as part of wounding |
| $C_{10}^{Clot, d0}$ | 300.0 kPa | Assumed in line with Ref. [71] |
| $C_{10}^{d0, lb}$ | 2.5 kPa | Linear extrapolation from values at days 7 and 14 |
| $C_{10}^{d7, lb}$ | 1.2 kPa | Lower bound from 95% CI in Fig. 3a |
| $C_{10}^{d14, lb}$ | 0.0 kPa | Lower bound from 95% CI in Fig. 3a |
| $k_1^{d0, lb}$ | 0.0 MPa | = $k_1^{i.w.}$ since no collagen is deposited prior to day 0 |
| $k_1^{d7, lb}$ | 0.6 MPa | Lower bound from 95% CI in Fig. 3b |
| $k_1^{d14, lb}$ | 1.2 MPa | Lower bound from 95% CI in Fig. 3b |
| $C_{10}^{d0, m}$ | 51.2 kPa | Linear extrapolation from values at days 7 and 14 |
| $C_{10}^{d7, m}$ | 25.9 kPa | Median value from Fig. 3a |
| $C_{10}^{d14, m}$ | 0.6 kPa | Median value from Fig. 3a |
| $k_1^{d0, m}$ | 0.0 MPa | = $k_1^{i.w.}$ since no collagen is deposited prior to day 0 |
| $k_1^{d7, m}$ | 19.0 MPa | Median value from Fig. 3b |
| $k_1^{d14, m}$ | 9.0 MPa | Median value from Fig. 3b |
| $C_{10}^{d0, ub}$ | 167.2 kPa | Linear extrapolation from values at days 7 and 14 |
| $C_{10}^{d7, ub}$ | 94.3 kPa | Upper bound from 95% CI in Fig. 3a |
| $C_{10}^{d14, ub}$ | 21.4 kPa | Upper bound from 95% CI in Fig. 3a |
| $k_1^{d0, ub}$ | 0.0 MPa | = $k_1^{i.w.}$ since no collagen is deposited prior to day 0 |
| $k_1^{d7, ub}$ | 86.2 MPa | Upper bound from 95% CI in Fig. 3b |
| $k_1^{d14, ub}$ | 148.1 MPa | Upper bound from 95% CI in Fig. 3b |

## Acknowledgments

This work was supported by the Swiss National Science Foundation (Doc.Mobility P1EZP2-178376) and by the US National Science Foundation (NSF-CMMI 1911346).

